# Photoclick Phase-separating Hydrogels for 3D Cell Culture and Volumetric Bioprinting

**DOI:** 10.1101/2022.01.29.478338

**Authors:** Monica Z. Müller, Margherita Bernero, Wanwan Qiu, Robert W. Style, Ralph Müller, Xiao-Hua Qin

## Abstract

Macroporous scaffolds facilitate solute transport and cell-cell communication, but materials allowing for *in situ* pore formation and 3D printing in aqueous solutions are scarce. Here, we introduce an efficient thiol-ene photoclick resin for light-assisted fabrication of cell-compatible macroporous hydrogels via photopolymerization-induced phase separation (PIPS). This resin consists of norbornene-functionalized polyvinyl alcohol, di-thiol crosslinker and dextran sulfate, which can rapidly form a hydrogel with interconnected pores by PIPS. The pore size is tunable in the range of 2-40 μm as a function of light intensity, polymer composition and molecular charge. Unlike conventional methods to porous materials, PIPS uniquely allows *in situ* pore formation in the presence of living cells, thereby enabling 3D cell culture and bioprinting applications. We demonstrate fast 3D photoencapsulation of living cells, enhanced cell spreading in macroporous hydrogels, and tomographic volumetric bioprinting of cm-scale hydrogel constructs with hierarchical pores within 20 seconds. Collectively, this resin is cell-compatible, low-cost, easy-to-make and highly efficient for PIPS, offering promises for fast photofabrication of living tissues with complex porous structures.

## 1. Introduction

Macroporous scaffolds^1-3^ have emerged as promising materials for 3D cell culture and tissue engineering since they facilitate solute transport, cell-cell communication and tissue growth. Methods to create macroporosity include porogen leaching^1-3^, microgel annealing^4-7^, microstrand molding^8^, and phase separation^9-13^. However, each approach has its respective limitations. Huebsch et al. created macroporous alginate hydrogels by adding hydrolytically degradable porogens for recruiting or releasing cells.^2^ Since hydrolytic degradation is a slow process, the embedded cells need at least 7 days to colonize and interact with the void space. Another method is phase separation, which avoids harsh treatments and offers better cell-compatibility. For instance, Broguiere et al. prepared macroporous polyethylene glycol (PEG) hydrogels via Michael addition for 3D culture of neurons.^9^ However, the required crosslinking takes ca. 1 hour long and the materials are not 3D printable, making them unsuitable for the fabrication of complex tissue constructs.

A major challenge in additive (bio)manufacturing^14-17^ is the shortage of bioresin formulations that allow digital fabrication of hydrogel constructs with both macroscopically and microscopically controlled architecture to support 3D cell growth and tissue regeneration. Existing hydrogel systems^15,18^ for 3D bioprinting often have very small pore sizes (5 nm - 50 nm) that fail to provide a permissive environment for encapsulated cells. A combination of 3D printing and phase separation has been explored in other materials than hydrogels such as glass^19^ and acrylic monomers^20,21^ using digital light processing^19,20^ and two-photon polymerization^21^. However, these methods rely on the removal of porogens by vacuum drying.

Liquid–liquid phase separation has raised increasing attention in life sciences owing to its vital role in human health and diseases^22,23^. Phase separation in living systems relies on macromolecules. Unlike small molecules, macromolecules tend to separate and form droplets in aqueous solutions when the mixing entropy is not favourable due to large molar mass^24,25^. Herein, we report fast construction of cell-compatible macroporous hydrogels by leveraging a thiol-ene photoclick resin^16,26^ and photopolymerization-induced phase separation (PIPS, Fig. 1a). In PIPS, photopolymerization induces phase separation, driven by changes in entropy as the molecular mass of polymers increases during *in situ* photocrosslinking. This process can be used to form porous microstructures within a hydrogel (Fig. 1b-c). We employ PIPS to produce cell-compatible macroporous hydrogels within seconds via efficient thiol-ene photoclick polymerization (Fig. 1d-e). Before photocuring, the resin is optically transparent, which is the key for light-based 3D printing such as tomographic volumetric bioprinting^17,27,28^. After PIPS, the materials form stable macroporous hydrogels in the presence of living cells.

**Fig. 1:**
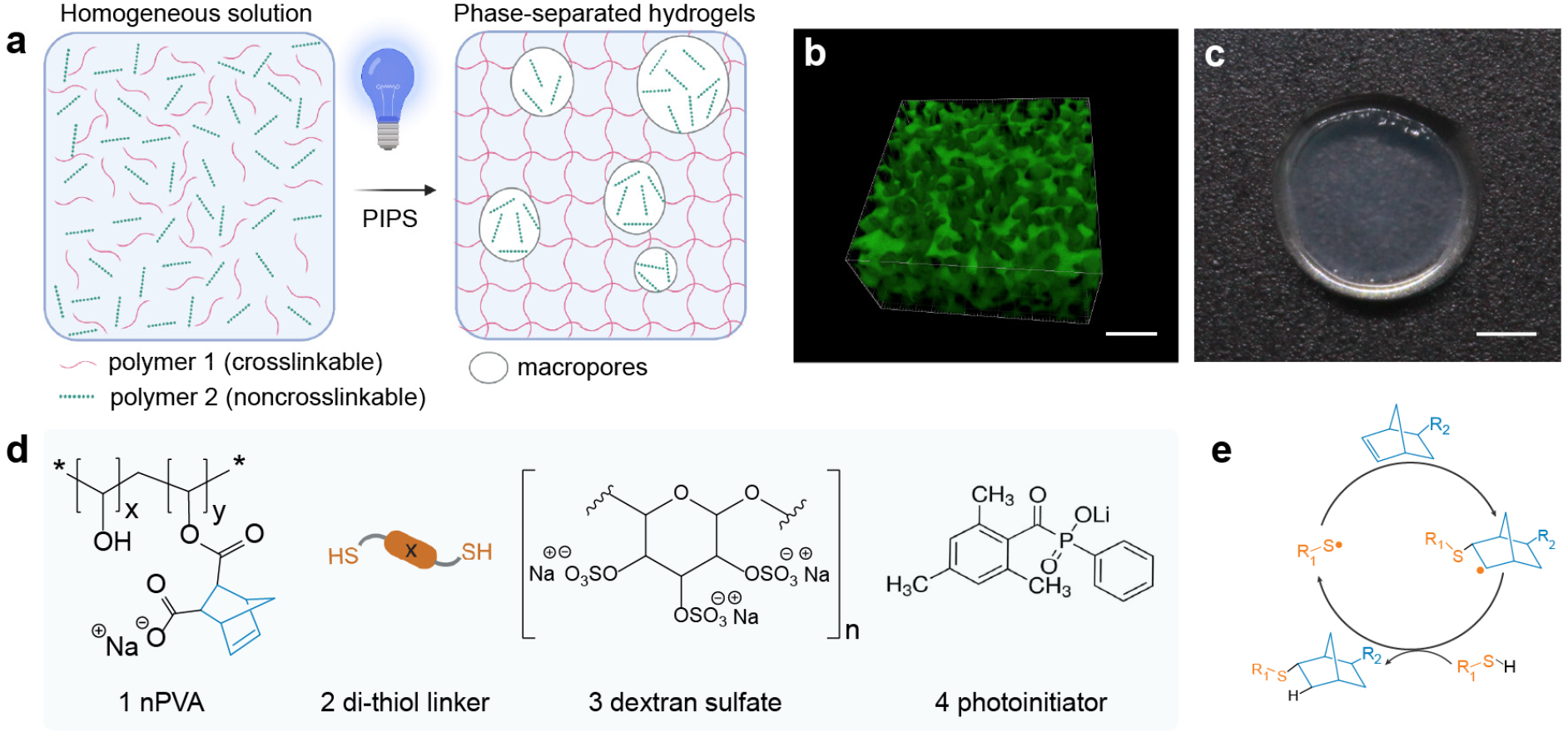
Macroporous hydrogels by photopolymerization-induced phase separation (PIPS). **a** Schematic representation of the PIPS process: two initially miscible polymers become immiscible during the photocrosslinking of one polymer because of entropy change, which induces phase separation and pore formation. **b** Representative confocal microscopy image of macroporous gels permeabilized with FITC-dextran. Scale bar = 10 µm. **c** Photograph of phase-separated hydrogels. Scale bar = 2 mm. **d** Chemical structures of key components for PIPS: 1) norbornene-functionalized polyvinyl alcohol (nPVA), 2) PEG-di-thiol (PEG-2-SH) linker, 3) dextran sulfate and 4) water-soluble photoinitiator (LAP, lithium phenyl-2,4,6-trimethylbenzoylphosphinate). **e** Schematic of the proposed mechanism in radical-mediated thiol-norbornene photoclick polymerization.

## 2. Results & Discussion

### 2.1 Design Considerations of a Phase-separating Resin

In the present study, we devised an efficient photoclick phase-separating resin based on the following considerations. First, we focused on the properties of aqueous mixtures of two nonionic polymers: polyvinyl alcohol (PVA) and dextran. These two polymers are known to phase separate as a function of temperature and salt additives because they have distinctive capacity to bind water molecules^29^. However, less is known about the phase-separating phenomena between ionic polymers. Second, we chose to develop a photoclick phase-separating resin made of synthetic polymers that are miscible before photocrosslinking but undergoes *in situ* pore formation upon exposure to light. We previously reported a synthetic polymer based on photoclick norbornene-functionalized PVA (nPVA)^26^, which is water soluble and highly efficient for hydrogel formation in the presence of thiols via step-growth thiol-ene photopolymerization. This process can be spatiotemporally controlled by light using a water-soluble photoinitiator such as LAP (Fig. 1d)^30^. Thus, we reasoned that aqueous mixtures of nPVA and an ionic derivative of dextran may undergo PIPS to form macroporous hydrogels in a spatially and temporally controlled fashion. We selected dextran sulfate (DS) as non-crosslinkable polymer due to its ionic nature and commercial availability. Third, the resin formulations must be cell-friendly to enable *in situ* photoencapsulation of living cells within hydrogels by means of state-of-the-art 3D cell culture and bioprinting techniques.

### 2.2 Evidence of PIPS

To determine potential compositions for PIPS, an approximate phase diagram of nPVA (*M*_*W*_, 61 kDa, degree of functionalization: 7%) and DS (*M*_*W*_, 40 kDa) was made (Fig. 2a and Supplementary Fig. 1). It resembles a phase diagram of PVA and dextran reported in literature^29^. Compositions were chosen between 2-3% nPVA and 2.5-3.5% DS, as the critical point (CP) was estimated to be within this region. We performed *in situ* photo-rheometry to evaluate the crosslinking of the compositions. The results showed rapid crosslinking within 30 seconds, as indicated by the sharp increase of the storage modulus (G’) upon exposure to light (Fig. 2b, Table 1).

**Table 1:** Composition and properties of common mixes used in this study. 0.05% LAP, PEG-2-SH crosslinker for a thiol:ene ratio at 4:5.

| Mix No. | nPVA % (w/w)<sup>a</sup> | DS % (w/w) | Pore size [$\mu\text{m}$]<sup>b</sup> | G' [Pa]<sup>f</sup> | G'' [Pa]<sup>f</sup> | G''/G'<sup>f</sup> |
| --- | --- | --- | --- | --- | --- | --- |
| 0 | 2.0 | - | NA<sup>d</sup> | 62.5 ± 2.0 | 1.1 ± 0.07 | 0.018 ± 0.001 |
| 1 | 2.0 | 2.5 | 3 | 111.4 ± 5.0 | 0.68 ± 0.12 | 0.006 ± 0.001 |
| 2 | 2.0 | 3.0 | 7 | 162.2 ± 3.6 | 0.75 ± 0.03 | 0.005 ± 0.0002 |
| 3<sup>c</sup> | 2.0 | 1.0 | 18 | 4249 ± 1063<sup>e</sup> | 94.2 ± 38.9<sup>e</sup> | 0.022 ± 0.004<sup>e</sup> |

<sup>a</sup> Weight concentration was used to correlate with the approximated phase diagram;
<sup>b</sup> determined by RIFFT of confocal imaging data after gel swelling;
<sup>c</sup> 5% gelatin was added to enhance the viscosity of resin;
<sup>d</sup> no pores detected;
<sup>e</sup> before incubation at 37 °C (prior to gelatin release).
<sup>f</sup> G', G'' and G''/G' are the plateau values after 5 min of UV irradiation.

**Fig. 2:**
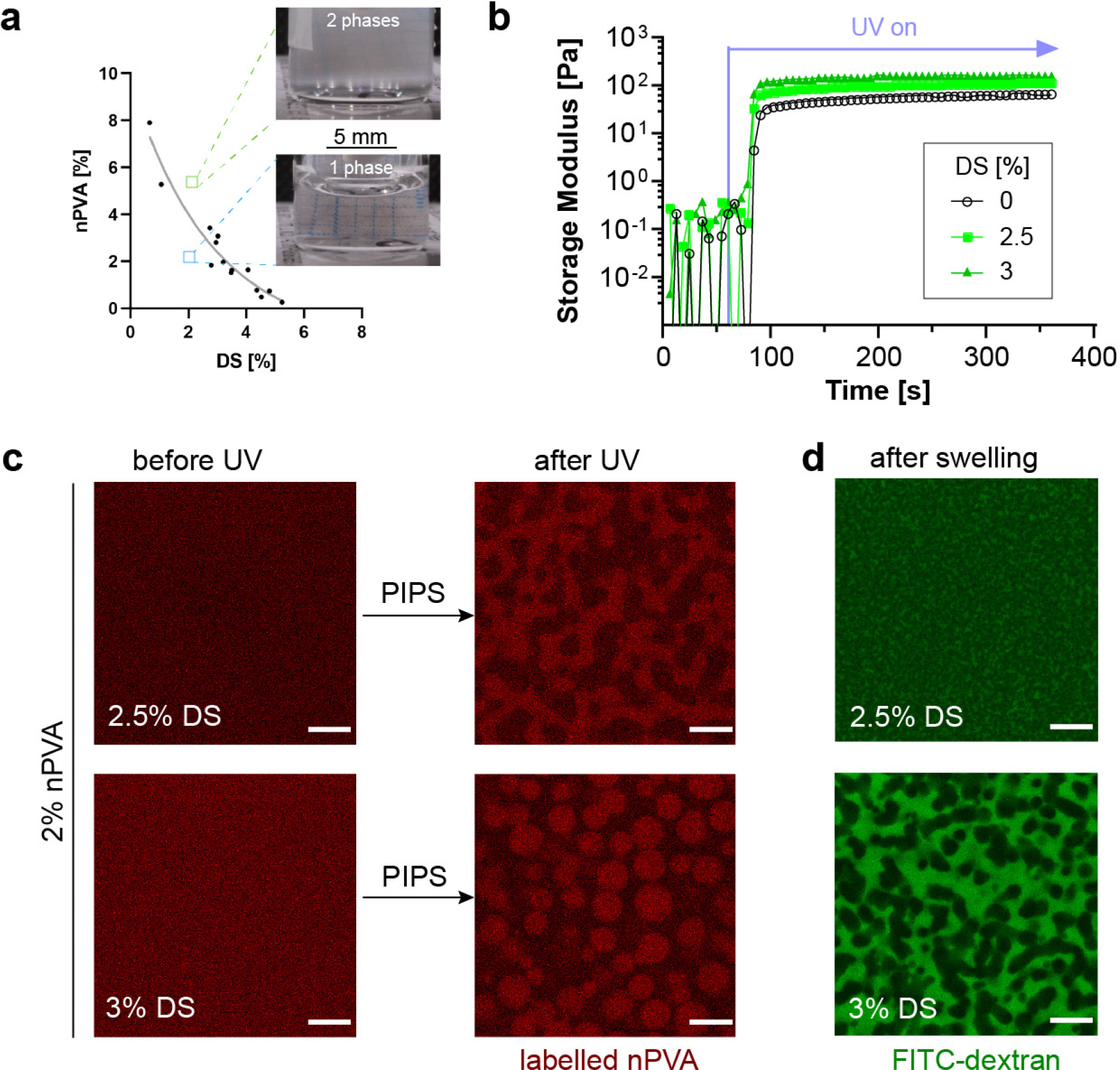
Macroporous hydrogels by PIPS between norbornene-functionalized PVA (nPVA) and dextran sulfates (DS). **a** Approximated phase diagram between nPVA (*M*_*W*_, 61 kDa) and DS (*M*_*W*_, 40 kDa). **b** Time sweep of storage (G’) and loss (G’’) modulus of 2% nPVA with 0, 2.5 and 3% DS) under UV irradiation after 60 s. **c** Evidence of PIPS. Representative confocal images of Mix 1 (2.5% DS) and Mix 2 (3% DS) before and after photocrosslinking. 1/50th of FITC-labelled nPVA (red) was used for confocal imaging. **d** Diffusion of FITC-dextran molecules into the pores. Representative confocal images of Mix 1 and Mix 2 crosslinked under UV irradiation after 3 days of swelling in PBS and then 2 days in a FITC-dextran solution (green, indicating the void space). Images from the middle of gels show that the dye has penetrated the matrix. Scale bars: 10 μm (**c, d**).

To investigate if the resin compositions undergo PIPS, FITC-labelled nPVA was added to the resins as a fluorescent reporter of phase separation. The samples before and after UV curing were investigated by confocal microscopy. As shown in Fig. 2c, porous structures were observed only after UV curing. This observation confirms that the gel precursors were initially miscible, and phase separated due to photocrosslinking. Moreover, the pore morphology can be tuned by the concentration of DS: 2.5% DS results in bi-continuous structures, while 3% DS results in droplets. The radial intensity plots of the Fast Fourier Transformation (RIFFT) of the images after curing show a clear peak, implying a uniform length scale (Supplementary Fig. 2a). Both peaks correspond to a length scale of around 8 μm. No phase separation was found in the samples with lower DS concentrations and without DS (Supplementary Fig. 3).

Since the pore size may change dramatically after swelling, a dye diffusion experiment was performed to verify the gel microstructure. Confocal microscope imaging data showed that FITC-dextran (*M*_*W*_, 500 kDa) could permeate the pore space (Fig. 2d, Supplementary Movie 1). The RIFFT showed a peak for both compositions (Supplementary Fig. 2b). No pores were seen in the sample without DS (Supplementary Fig. 4). The sample with 3% DS had a length scale of around 8 μm. Surprisingly, the length scale for 2.5% DS was found to be around 3 μm, which is smaller than the length scale observed before swelling. These results suggest that the pores are permeable by FITC-dextran tracers and the pore size can be tuned by varying the DS concentration.

### 2.3 The Impact of Resin Composition on PIPS: Pore Size, Gel Mechanics and Turbidity

To study the effect of molecular charge on PIPS, the phase-separating resin was compared to a composition with the nonionic dextran instead of DS. With the same concentration, no pores were seen in the sample with dextran while pores were seen with DS (Fig. 3a). RIFFT analysis revealed that there is no peak in the sample with dextran, whereas a peak corresponding to a pore size of 5 μm was found in the sample with DS. These results imply that charged polymers promote pore formation and highlight the uniqueness of negatively charged DS for PIPS, which may repel the negatively charged nPVA more efficiently than dextran of equal *M*_*W*_.

**Fig. 3:**
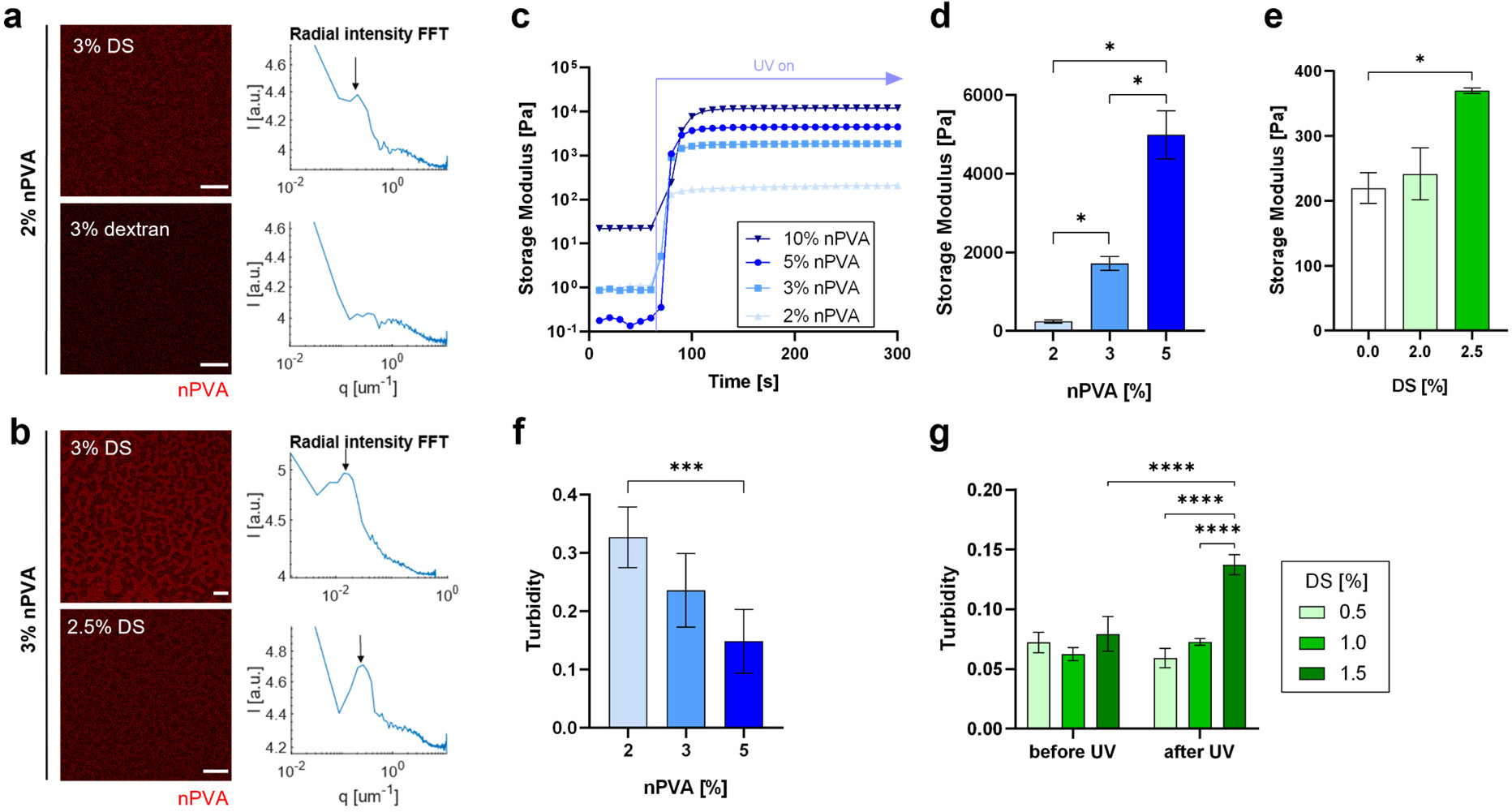
The effect of material composition on structural and physical properties of PVA hydrogels. **a** Molecular charge promotes the phase separation between nPVA and DS. Representative confocal images (left) of the composition containing 2% nPVA and 3% DS after UV curing. FITC-labelled nPVA (red) was used for confocal imaging. RIFFT of the images showing a clear peak with DS. **b** Phase separation with 3% nPVA and different DS concentrations resulting in pores at different scales. Representative confocal images after UV curing and RIFFT showing clear peaks (arrows). **c-e** In situ photo-rheometry of hydrogels with varying nPVA (**c, d**) and DS (**e**) concentrations; 10 rad s^-1^ and 0.5% strain. **c** Representative time sweep of G’ of gels with 2% DS and varying nPVA under UV after 60 s. **d-e** G’ of gels with either 2% DS (**d**) or 2% nPVA (**e**) after 4 min of crosslinking (N=3); p-values: p=0.0102 (2 vs. 3% nPVA), p=0.0115 (2% vs. 5% nPVA), p=0.0258 (3% vs. 5% nPVA), p= 0.0177 (0% vs. 2.5% DS). **f-g** Gel turbidity is higher in phase-separated compositions close to the critical point (N=5). **f** Turbidity of cured 2% nPVA gels with 2.5% DS showing significantly higher turbidity than 5% nPVA; ***: p= 0.0009. **g** Turbidity of 4% nPVA gels with 0.5, 1 and 1.5% DS before and after UV curing; ****: p <0.0001. Scale bars: 10 µm (**a, b** bottom image) and 50 µm (**b** top image). Data presented as mean ± SD.

Resins with a higher nPVA content were explored as alternative compositions. A resin containing 3% nPVA and 3% DS exhibited larger pores after PIPS (Fig. 3b) with the peak of the RIFFT corresponding to a length scale of 35 μm. With 2.5% DS, pores on a length scale of 4 μm were observed. At 3.5% DS, the composition phase separated even before gel formation (Supplementary Fig. 5). Compositions with higher nPVA concentrations were also screened, but the pores were either not interconnected or less homogenously distributed (Supplementary Fig. 6), presumably due to phase separation before photopolymerization.

Photo-rheology was used to determine the physical properties of these hydrogels as a function of different compositions. The G’ was higher with increase of nPVA with the same thiol-ene ratio (Fig. 3c-d). The mean values were 146, 1’344 and 3’186 Pa for 2, 3 and 5% nPVA, respectively. These data indicate the increase of crosslinking density as the polymer content increases. Notably, the photo-curing of a resin with 1% nPVA and 2.5% DS was unsuccessful due to insufficient crosslinking at low polymer content.

Next, we tested if the variation of DS content leads to changes in gel stiffness (defined as the G’-plateau). Interestingly, the stiffness was significantly higher for the sample with 2.5% DS compared to the control without DS (Fig. 3e). This is likely due to the effect of partitioning where nPVA and PEG-2-SH crosslinker may be rich in one phase whereas DS is rich in the other phase. This may lead to a higher crosslinking density in the nPVA-rich phase, resulting in a higher gel stiffness. The amplitude and frequency sweeps showed that a strain of 0.5% was within the linear viscoelastic region and the gels exhibited a solid behavior over a wide range of frequencies (Supplementary Fig. 7). Interestingly, the mass swelling ratio of 2% nPVA hydrogels with varying DS content showed no significant difference (Supplementary Fig. 8). Future work is needed to investigate the dynamics of solute transport and swelling in these macroporous hydrogels.

We studied the influence of gel composition on turbidity of the hydrogels on a plate reader. The turbidity was higher in hydrogels with compositions close to the CP. For instance, hydrogels with 2% nPVA and 2.5% DS showed a significantly higher level of turbidity compared to 5% nPVA gels with the same DS concentration (Fig. 3f). Additionally, the turbidity of formulations with 4% nPVA and varying DS before and after UV curing was measured. Before UV curing, there was no significant difference in the turbidity. After UV curing, the turbidity of hydrogels with 1.5% DS was significantly higher compared to 0.5% DS and 1% DS groups as well as all groups before UV curing (Fig. 3g). This indicates that the composition with 4% nPVA and 1.5% DS was initially miscible, but phase separated upon photopolymerization.

### 2.4 The Impact of Light Intensity on PIPS

Tuning the pore size by light intensity will significantly expand the applicability of phase-separating resins for light-based 3D printing. Such tuning has been only shown in purely organic polymeric matrices in literature.^31^ Thus, we investigated the effect of light intensity (5 – 100 mW cm^-2^) on PIPS and pore size distribution. As anticipated, imaging of gels cured at different light intensities showed that the length scale of the pores can be tuned by the UV light intensity (Fig. 4a). For irradiation at an intensity of 5 mW cm^-2^, the pore size was 8 μm. As the light intensity increased to 10 mW cm^-2^, the pore size decreased to 5 μm. Further, it decreased to 2 μm when the light intensity increased to 100 mW cm^-2^ (Fig. 4b). The best fit was obtained with the following function:

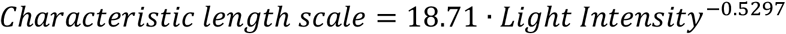

**Fig. 4:**
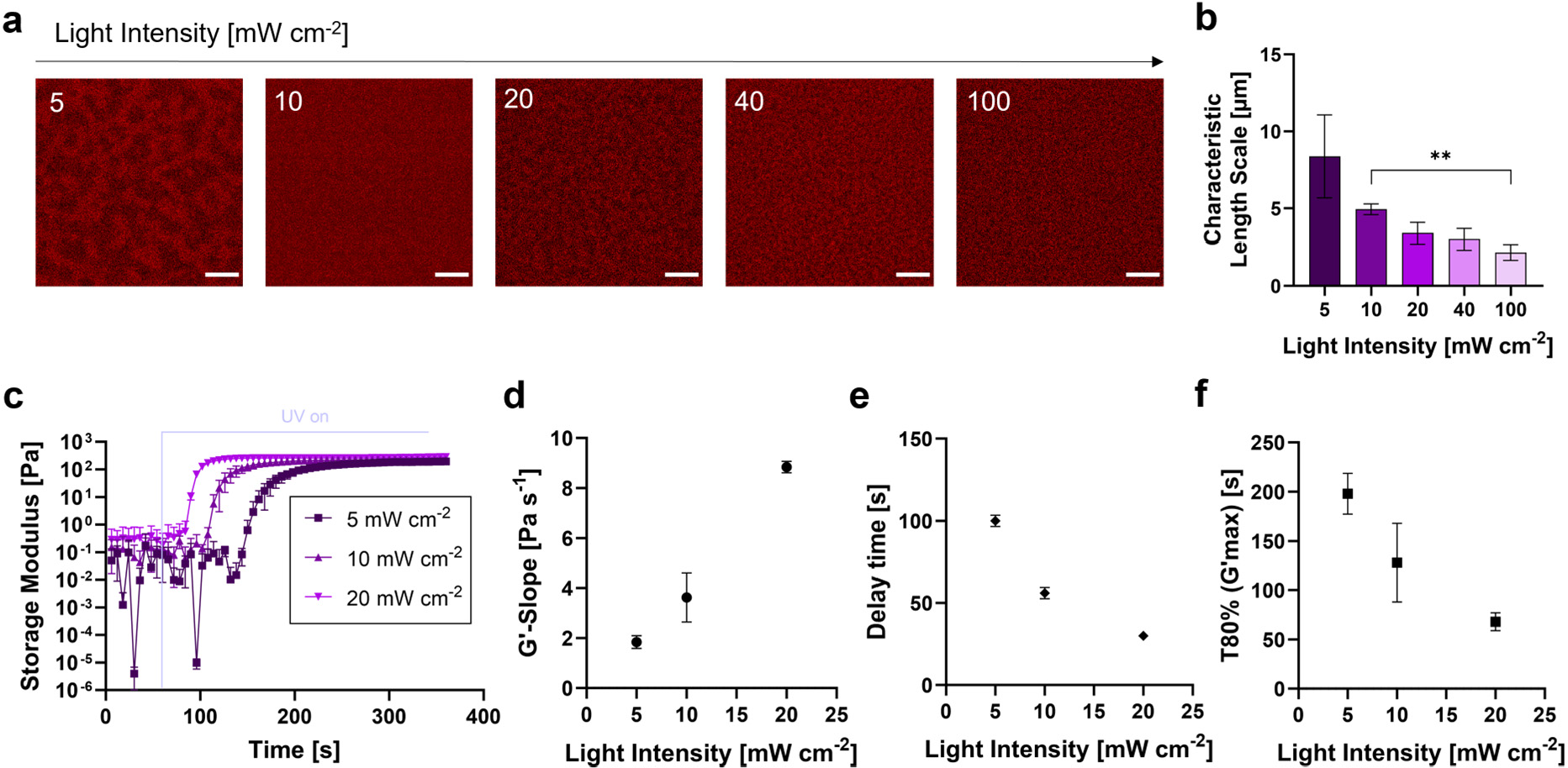
The effect of light intensity on pore size and photocrosslinking dynamics. **a-b** Characteristic length scale of the pores can be tuned as a function of the light intensity. **a** Representative confocal images of UV-cured hydrogels (2%nPVA-3%DS) with light intensities ranging from 5 to 100 mW cm^-2^ (365 nm). **b** Characteristic length scale obtained from RIFFT plot peaks of such images (N=3); curve fit with a power function. **c-f** Crosslinking dynamics with different light intensities: effect on the time sweep of gel storage moduli (**c**) and G’-slope (**d**), delay time for crosslinking (**e**) and time to reach 80% of the G’-plateau (**f**) under UV irradiation from 60 s at different light intensities (5-20 mW cm^-2^). Data presented as mean ± SD. Scale bars = 10 μm (**a**).

Kimura et al. investigated the process of PIPS in a mixture of polystyrene and methylacrylate^32^. By changing the light intensity, they obtained a variety of stationary morphologies. With higher light intensity, the pore morphology has less time to evolve before it is arrested at the onset of the gelation.

Photo-rheometry measurements showed that an increase in light intensity accelerates crosslinking (Fig. 4c-f). The evolution of G’ with different UV light intensities is shown in Fig. 4c. For analyzing the rate of crosslinking, a linear regression at the G’ plots. The slope, which indicates the rate of crosslinking, was higher with increased light intensity (Fig. 4d). The delay time denotes the timepoint where the linear regression curve starts (Fig. 4e), whereas T_80%_ refers to the time to reach 80% of the maximum G’ within 5 min of UV curing. When increasing the light intensity from 5 mW cm^-2^ to 20 mW cm^-2^, the crosslinking speed is increased from 1.8 Pa s^-1^ to 8.8 Pa s^-1^, whereas the delay time is decreased from 100 s to 30 s, and the time to reach T_80%_ is decreased from 198 s to 68 s, respectively. The results confirmed that photocrosslinking is faster at higher light intensities (Fig. 4f). Together, these findings suggest that the light intensity can be used to tune the crosslinking dynamics and PIPS, which influences the length scale of the pores.

### 2.5 3D Cell Culture

For studying cellular behavior in the phase-separating gels, human mesenchymal stem cells (hMSCs) were encapsulated in the hydrogel at a density of 0.5 × 10^6^ cells/mL. The cells were mixed with the precursors and cured under UV irradiation (Fig. 5a). All compositions showed a significantly high cell viability (>90%) (Fig. 5b-c). The phase-separating compositions showed a higher average cell area compared to the control. Cells in 3% DS gels showed a significantly higher cell area compared to the control without DS on day 13 (Fig. 5c). Further, the cell area significantly increased over time. Similarly, confocal images of actin-nuclei stained cells showed longer processes in the phase-separating gels (Fig. 5c, Supplementary Movie 2) after 13 days osteogenic culture. The length of cell processes increases over time in the phase-separating compositions (Supplementary Fig. 9). Cells penetrating through the pores were also observed (Supplementary Fig. 10).

**Fig. 5:**
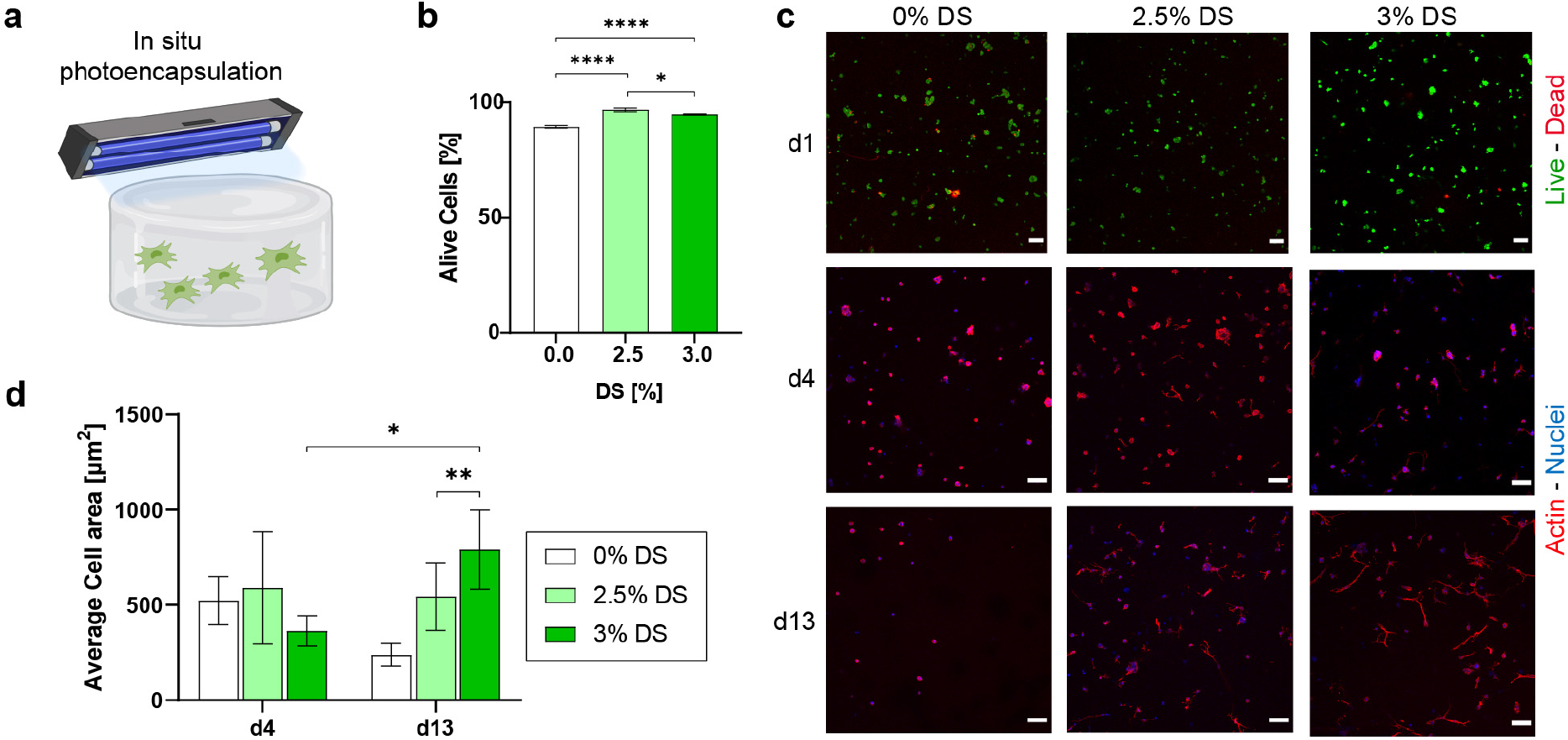
3D Photoencapsulation of hMSCs in macroporous PVA hydrogels. **a** Schematic representation of *in situ* cell photoencapsulation, where the gel is crosslinked under UV in the presence of live hMSC cells. **b** Cell viability on day 1 as determined by a live-dead assay (N=3); one-way ANOVA with Tukey’s correction for multiple comparisons, *: p = 0.0180, ****: p < 0.0001. **c** Representative confocal images of live-dead stained hMSCs on day 1 and actin-nuclei-stained cells on day 1, 4 and 13. Maximum intensity projections (MIP) from z-stacks (100 μm). Scale bars = 100 μm. **d** Average cell area after 4 and 13 days of 3D osteogenic culture (N=3, 50 μm thick z-stacks); two-way ANOVA with correction for multiple comparisons with Sidak for comparison of the two timepoints and with Tukey for comparisons of the three compositions, *: p = 0.0358, **: p = 0.0063. Data presented as mean ± SD.

### 2.6 Volumetric Bioprinting with An Optimized Resin

Volumetric bioprinting (VBP)^16,17,27,33^ is an enabling technique which allows rapid construction of 3D living hydrogel constructs in a rotating glass vial in a single step, addressing the limitations of conventional layer-by-layer manufacturing. The phase-separating resins were combined with the Readily3D tomographic volumetric printer (Fig. 6a). Initial printing attempts were unsuccessful due to insufficient viscosity of the resins. In VBP, a viscous or physically crosslinked resin is often needed to eliminate or limit sedimentation of structures during printing. To meet this requirement, we added 4-5% sacrificial gelatin to the resin as a viscosity enhancer. A resin containing 2% nPVA, 1% DS, and 5% gelatin (Mix 3, Table 1) showed homogenous distribution of porosity. Unwanted phase separation was seen with 1.5% DS, while no phase separation was seen with 0.5% DS (Supplementary Fig. 11). With 1% DS, pores in the range of about 2-5 μm were observed (Supplementary Fig. 12).

**Fig. 6:**
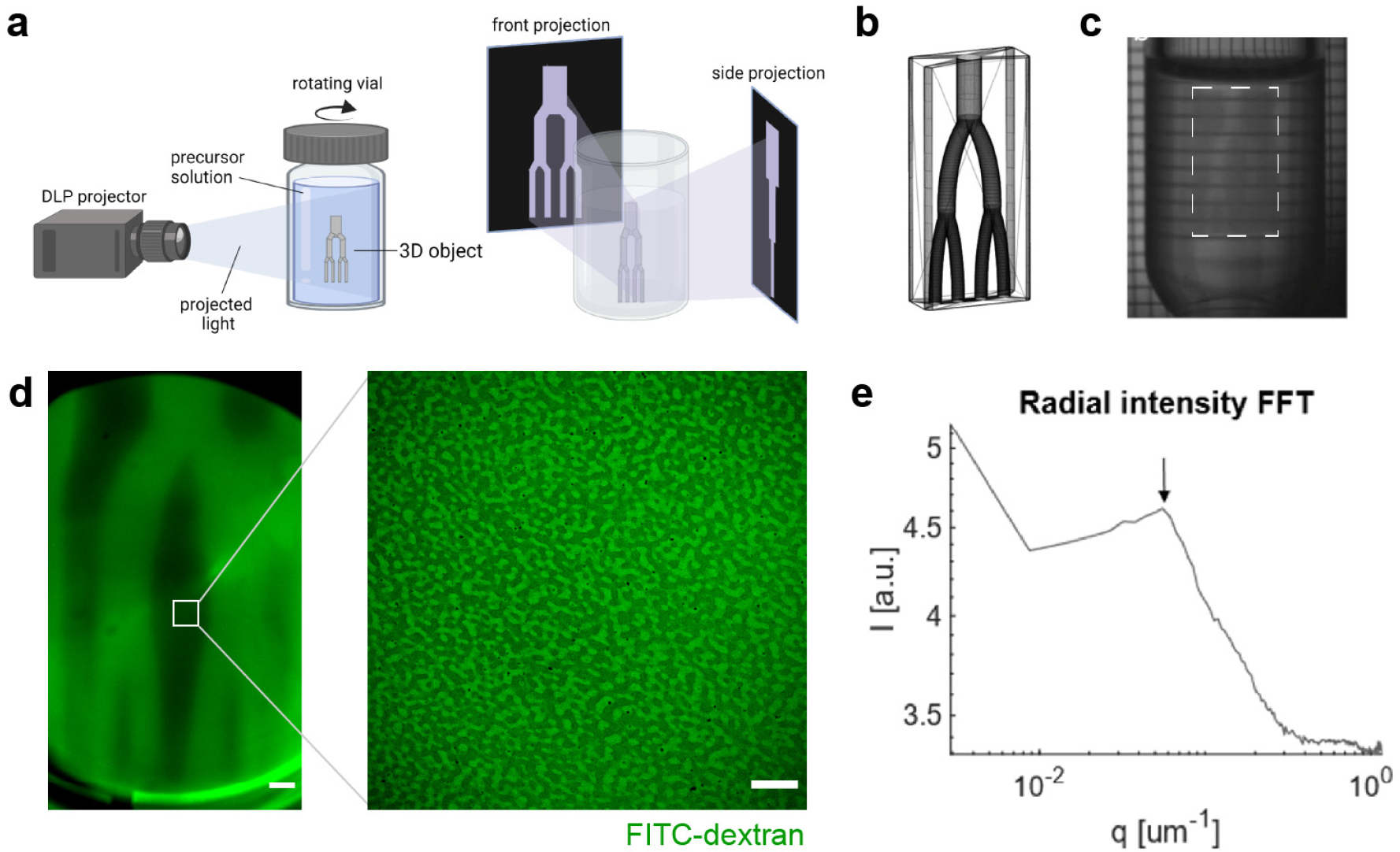
Tomographic volumetric printing of macroporous hydrogels using an optimized phase-separating resin. **a** Schematic representation of the volumetric printing process. **b** STL model of a vascularized branch model. **c** Photograph of the sample during the volumetric printing process with the appearing 3D object marked by a rectangle. **d** Representative confocal image at the center of the construct after two days in FITC-dextran. **e** RIFFT of the image shown in **d** (right) showing a clear peak (arrow) indicating a uniform length scale of 18 μm for the porosity. Scale bars: 1 mm (**d**, left) and 50 µm (**e**, right).

Further, the resin exhibited temperature-dependent gelation. A temperature sweep from 37 °C to 8 °C showed a gelation point at around 20-22 °C (Supplementary Fig. 13a). Above 22 °C, the complex viscosity was about 100 mPa·s (Fig. 13b), which is almost seven times higher than the composition without gelatin. Crosslinking the composition at 25 °C is indicated by a clear increase in G’ and G’’ after UV irradiation (Supplementary Fig. 13c) until reaching a plateau in G’ of about 2 kPa was reached within less than one minute. Laser dose tests were performed to determine a suitable laser dose range for the resin. The volumetric prints and laser dose tests were conducted above room temperature to ensure the resin remaining as a viscous liquid. A test with small intervals was performed and a threshold of 75 mJ cm^-2^ was found (Supplementary Table 1). 3D objects could be printed with a laser dose in the range of 95 to 125 mJ cm^-2^.

Using volumetric printing, a vascular branch model was printed at a laser dose of 95 mJ cm^-2^ within 12 seconds (Supplementary Movie 3). The computer-aided design (CAD) model (Fig. 6b) has a height of 1.4 cm, vascular channels with dimensions of 0.7-1.4 mm. The turbid construct is depicted while printing (Fig. 6c). Channels were shown to be perfusable after leaching out the gelatin for two days at 37 °C and subsequent incubation in a FITC-dextran solution for one day (Fig. 6d). Next, we investigated if the printed constructs are indeed macroporous. After incubation in PBS and FITC-dextran, confocal imaging data showed interconnected pores in the printed gel constructs (Fig. 6d). The RIFFT of this image showed a clear peak corresponding to characteristic length scale of around 18 μm (Fig. 6e).

An initial experiment showed that the branch structure could be printed in the presence of live hMSCs. After bioprinting, the channels were perfused by phenol red stained medium (Supplementary Fig. 14a). The gel appeared to be macroporous (Supplementary Fig. 14b). Cells appeared highly viable as evidenced by a live staining, but no spreading was observed over the culture period. Therefore, the bioresin formulation was supplemented with CGRGDS peptide to promote cell adhesion. We investigated the influence of DS content on resin stability at different temperatures. Large bubbles remained visible for 0.5% DS gels after crosslinking and only 0.2% DS gels showed a homogeneous porosity (Supplementary Fig. 15a). To reduce turbidity when printing with cells, the gelatin concentration was decreased to 4%. Thus, the bioresin composition contained 2% nPVA, 0.2% DS and 4% gelatin, and produced a macroporous hydrogel with a pore length scale of 2.2 µm (Supplementary Fig. 15b,c).

Additionally, optical tuning of the bioresin’s refractive index was investigated in accordance with a report by Bernal et al.^34^. By more closely matching the refractive index of cellular components, light scattering at the interface is reduced which enables VBP with improved resolution (Fig. 7a,b). The supplementation of our bioresin with 10% iodixanol successfully reduced the solution’s turbidity while printing (Supplementary Movie 4). This resulted in an increased cell viability on day 2 compared to a resin without iodixanol (93% and 62% respectively, Fig. 7c,d). Actin-nuclei staining indicated cells in the iodixanol-containing constructs spread (Fig. 7e), and showed a significantly increased cell area after 14 days in osteogenic culture (Fig. 7f). Alizarin Red S staining of the constructs indicated increased matrix mineralization for both groups after 14 days (Supplementary Fig. 16).

**Fig. 7:**
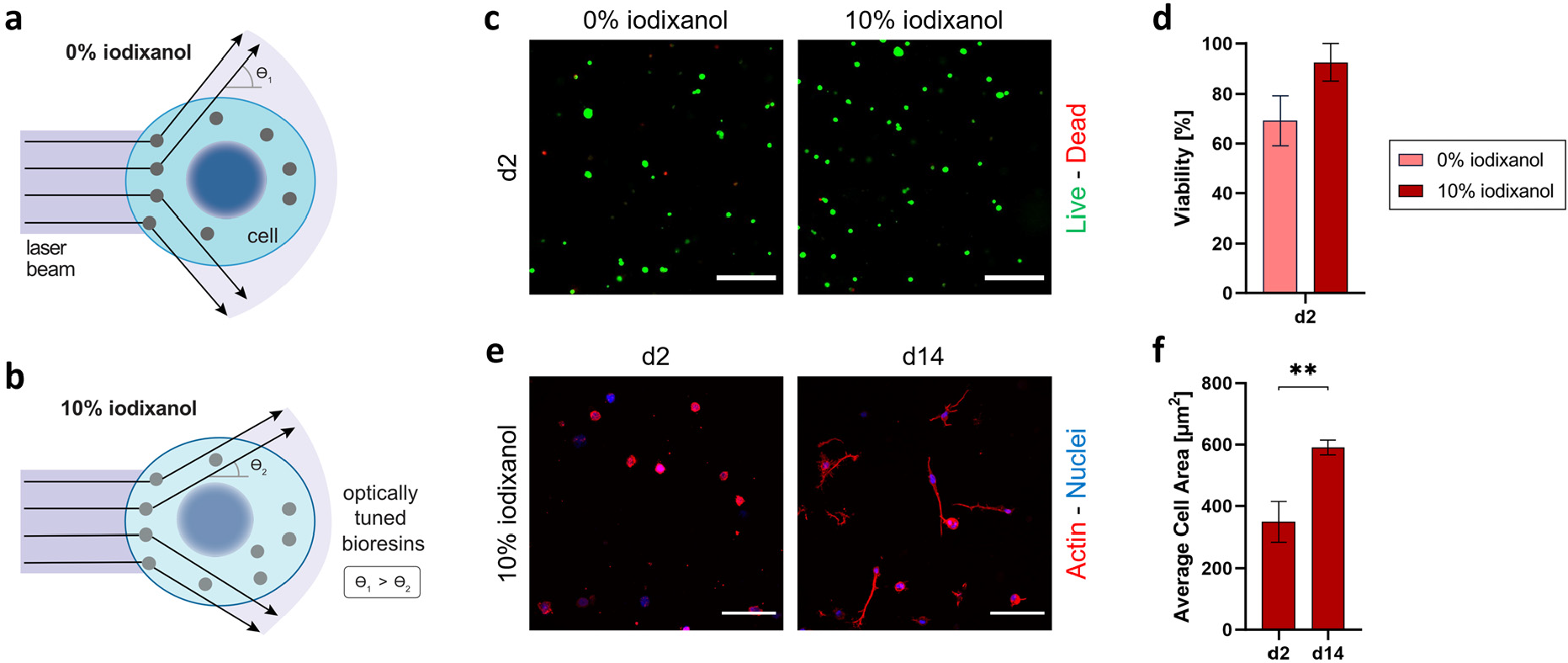
Volumetric bioprinting of a phase-separating resin with embedded hMSCs. **a-b** Schematic of the optical tuning mechanism visualizing reduced light scattering when comparing bioresins without (**a**) and with (**b**) 10% iodixanol. **c** Representative confocal maximum intensity projections (100 µm z-stacks) of live-dead stained hMSCs on day 2. Scale bars = 200 µm. **d** Cell viability quantification by a live-dead assay (N=3); Unpaired t test, non-significant difference. **e** Representative maximum intensity projections (100 µm z-stacks) of actin-nuclei-stained hMSCs in constructs with 10% iodixanol. Scale bars = 100 µm. **f** Average cell area quantification after 2 and 14 days of osteogenic culture for constructs printed with 10% iodixanol (N=3); Unpaired t test; p = 0.0039.

In conclusion, a photoclick phase-separating resin for fast construction of macroporous hydrogels was developed. The pore size could be fine-tuned by the gel composition as well as light intensity. The newly developed phase-separating nPVA gels showed a high viability on day one and increased spreading over two weeks. First successes were shown with volumetric bioprinting, where a cm-scale construct with macropores could be printed in under 20 s. To the best of our knowledge, the present study is the first report of PIPS for 3D printing of cell-compatible macroporous hydrogels. The developed phase-separating resins could be applicable to several fields of research. Besides VBP, the bioresin may find applications in other light-based 3D printing such as digital light processing (DLP)^35,36^, extrusion printing^37,38^ and two-photon direct laser writing^20,26,39^. Using light as the trigger, different pore sizes may be digitally 3D-printed within one object by tuning the light intensity and PIPS. Accordingly, complex hierarchical structures with macropores could be generated in the presence of living cells, which may significantly advance the fields of biofabrication and functional tissue engineering.

## 4. Methods

### Materials

A final concentration of 0.05% LAP was used throughout all experiments with UV curing. nPVA (61 kDa, DoF 7%) was synthesized as described elsewhere^40^. It was used at different concentrations and dissolved in a solution containing the photoinitiator by vortexing and ultrasonication. PEG-2-SH (2 kDa, LaysanBio) was used as a crosslinker with a concentration that results in a thiol/ene ratio of 0.8. It was dissolved freshly in PBS. DS (40 kDa, Carl Roth) was used for phase separation and was dissolved in PBS at 50°C for two hours by vortexing. Dextran (40 kDa, Sigma-Aldrich) was dissolved similarly. In cell culture experiments, 1 mM CGRGDSP (China Peptides) was added to the resins to promote cell attachment. All components were dissolved in PBS. Detailed description of resin compositions is included in the Supplementary.

### Approximation of a Phase Diagram

Vials with stock solution of 10% nPVA were prepared and then diluted with PBS, providing a set of vials with varied concentrations. These samples were titrated with 50% DS (40 kDa) in small steps until the cloud point. Another set of vials of differing starting concentrations was prepared and titrated with DS in decreasing steps, the last step before the cloud point was set to be 0.1 μL to increase accuracy. Careful observation of streaks appearing close to the cloud point helped in assessing how close each sample was to the cloud point.

### Hydrogel Preparation

The components were thoroughly mixed by pipetting up and down 20 times after each additional component. With large volumes (mL), the composition was mixed until it turned visibly clear, whereas it was mixed 20 times and then vortexed for two to three seconds with small volumes (μL). The crosslinker was usually added last. A detailed description of crosslinking and casting the gels in molds can be found in the supplementary methods.

### Permeability of Macroporous Hydrogels with FITC-Dextran

The permeability of gels with FITC-dextran (500 kDa, Sigma-Aldrich) was studied similar as described elsewhere^9^. In short, the gels were washed for three days at 4°C in PBS till reaching a swelling equilibrium. Then, PBS was replaced with a solution of FITC-dextran (0.1 mg/mL), and the samples were imaged after two days incubation.

### Turbidity Measurements

The turbidity was determined by measuring the absorption at 405 nm with a plate reader (Spark M10, Tecan). Before curing, 40 μL of the mixed precursor solution was loaded into a well of a 96 well plate. Gels cured in Teflon molds were punched with a 5 mm biopsy punch to fit into the plate and the turbidity was measured after adding 100 μL PBS into the well (N=5).

### Volumetric (Bio)-Printing

VBP was performed a Tomolite 1.0 bioprinter (Readily3D) together with the software Apparite. First, a laser dose test was conducted, in which defined spots were irradiated with varying laser intensity and irradiation time. After irradiation, it was noted which spots were polymerized. A second test with a smaller range was then conducted to determine the threshold needed for polymerization. For acellular printing, 3-4 mL of precursor solution was loaded into 18 mm diameter Pyrex vials. Bioprinting of the resin including the RGD-peptide was performed with 700 µL of resin in 10 mm diameter vials. A refractive index of 1.37 was chosen and the peak-to-average ratio was set to 6:1. The optimal settings as calculated by the software were used.

### 2D Fast Fourier Transform

2D-FFT is a common way to characterize the length scale of pores in bi-continuous structures^41^. The analysis was done with a MATLAB code, which can be found in the Supplementary. In short, an image is read and converted to greyscale. Then the 2D-FFT of the image multiplied by a hamming window is taken. Further, radial binning is performed to show the radial intensity of the FFT (RIFFT). The length scale was then calculated as q^-1^.

### Porosity Analysis

The threshold was set by Otsu’s method and the area fraction was measured. 3 images (N=3) in different locations of the same z-stack were analyzed. Analyses where the background signal was too high were not included in the results.

### Statistical Analysis

Statistical analysis was performed in GraphPad Prism 8.2.0, except for the slope analysis, which was performed in Excel. Ordinary one-way or two-way analysis of variance (ANOVA) were performed depending on the number of variables. They were followed by Sidak’s or Tukey’s test for multiple comparisons for two or more groups respectively. If the standard deviations did not appear equal among groups, the Brown-Forsynthe and Welch ANOVA test followed by Dunnett’s multiple comparison test was applied. In the two-way ANOVA, only simple effects within rows and columns are analyzed. Welch’s t test was used when there were only two groups. P values less than 0.05 were considered significant.

## Acknowledgements

This work was partly supported by the SNSF (no. 190345, 206501). We thank Esteban Oggianu and Dr. Paul Delrot (Readily3D) for experimental assistance and Alba Sicher for providing the MATLAB code. We also gratefully acknowledge staff members at the ETH Zurich ScopeM for their support and assistance in this work. W.Q. gratefully acknowledges financial support from China Scholarship Council (no. 202006790027). M. B. likes to acknowledge support from the ALIVE | Advanced Engineering with Living Materials initiative of ETH Zurich, which is funded by the ETH domain Strategic Focus Area Advanced Manufacturing programme.

## Author Contributions

Conceptualization: X.Q., M.M.; Experiments: M.M., M.B., W.Q., X.Q.; Data analysis: M.M., M.B., W.Q., R.S., R.M., X.Q.; Visualization: M.M., M.B., W.Q., X.Q.; Methodologies: M.M., M.B., W.Q., R.S., X.Q.; Writing of first draft: M.M., X.Q.; Reviewing of final manuscript: M.M., M.B., W.Q., R.S., R.M., X.Q.

