## Supplementary Materials for "Photoclick Phase-separating Hydrogels for 3D Cell Culture and Volumetric Bioprinting"

**Content:**

- Supplementary Methods
- Supplementary Figures 1-16 and Table 1-4
- MATLAB Script for FFT Analysis
- Supplementary Movies 1-4

**Supplementary Methods**

*Hydrogel Precursor Preparation*

In some experiments, a "10%" DS stock solution diluted by volume from a 50% stock solution was used. However, the density of the 50% stock solution was shown to be around 1.25 g/mL (Figure S17), so that the concentration of the diluted "10%" stock solution and so also the final DS concentrations in the hydrogel precursor mixture were higher than anticipated. The slightly higher average density of the diluted 10% stock solution compared to a directly prepared 10% stock solution also hints at this. In this report, 2.5% DS and 3% DS with 2% nPVA refer to compositions prepared from the diluted 10% DS stock solution, which correspond to actual concentrations of around 3.125% and 3.75% DS respectively as calculated by the density measurements.

*nPVA-DS Phase-Separating Composition*

The composition for Mix 1 is described in detail in **Table S2**. The components were added in the order listed, where it was especially important that the crosslinker is added last. For cell embedding, the order of mixing was altered to minimize the time where cells are outside of their medium (**Table S3**).

*The (Bio)resin Based on nPVA-DS-Gelatin for Volumetric Printing*

Gelatin was added because it gelates at lower temperatures, which makes it suitable to prevent sedimentation in volumetric printing. Gelatin was dissolved at 40 °C for 2-3 hours for a stock solution of 10% (w/v). The composition used for acellular printing experiments is described in **Table S4**. The components were mixed in this order and were warmed to prevent gelation during mixing. For volumetric bioprinting, the composition was supplemented with a CGRGDS peptide to promote cell adhesion as well as 100 ppm pyrogallol to prevent premature crosslinking. Iodixanol (OptiPrep, StemCell Technologies) was added to tune the resin's refractive index and reduce light scattering by cellular structures. Cells were resuspended in the gelatin portion for a final density of 2 x $10^6$ cells/mL. Composition details are described in **Table S5**.

*Gel Casting*

The gels were casted into PDMS molds with a thickness of 0.5 mm in wells with a diameter of 3 mm, which were punched out with biopsy punches. The molds were attached to a coverslip (30 mm Ø) or into a No. 1.5 glass bottom confocal dish (VWR). The wells were filled with 7 μL of precursor solution each. Sigmacoated coverslips were placed on top of the mold. For experiments with larger volumes, a Teflon mold was filled with 33 μL and covered with a

Sigmacoated-treated glass slide. After curing, PBS was added to the edge of the glass to facilitate loosening of the slide. The hydrogels were then transferred with a plastic spatula to a 12-well or 24-well plate filled with 1 mL of PBS for swelling.

The gels were cured for 5 min under UV lamps with an intensity of 18 - 20 mW cm$^{-2}$. The LAMAG UV lamp (365 nm) was used for most acellular experiments, the Navanino UV lamp (365) for most cell embedding experiments and the Thorlabs UV lamp (365 nm) for tuning the light intensity. After curing, the sample was directly imaged. Washed samples, however, were imaged after at least 4 days incubation in PBS at 4 °C. For the intensity tuning experiment, the gels with the light intensities 5 and 10 mW cm$^{-2}$ were cured for 20 and 10 min respectively to ensure complete crosslinking. A power meter (S120VC, Thorlabs) with a sensor (PM100D S12VC, Thorlabs) was used to measure the light intensities.

*Swelling Ratio*

Hydrogel discs (dimeter, 6 mm; height, 2 mm) were swollen in PBS at 37 °C for two days, the equilibrium wet mass ($M_w$) and the dried mass ($M_d$) of each gel disc were recorded, respectively. A sample size of 3-4 was used. The mass swelling ratio ($Q_m$) was calculated as:

$$Q_m = M_w/M_d$$

*Photo-Rheology*

In situ photo-rheological measurements were performed on a modular photo-rheometer (Anton Paar MCR 302) equipped with a 20 mm parallel plate geometry, a glass floor and UV-LED lamp (Thorlabs, Germany). The tests were carried out at room temperature (22 °C) unless otherwise noted. Mineral oil was loaded surrounding the sample to prevent from drying during testing. The time-sweep of shear storage and loss modulus were collected under irradiation of 365 nm UV-LED lamp (20 mW cm$^{-2}$) in triplicates at 0.5 % strain and 1 Hz frequency with 100 µm gap. Frequency and amplitude sweeps were performed to assess the mechanical behavior of the hydrogels after curing. Right after the UV crosslinking measurement, the frequency sweep was done *in situ* with an angular frequency of 0.1 – 100 rad/s and a strain of 0.5%. Amplitude sweeps were performed with an angular frequency of 10 rad/s and a strain of 0.01 – 100% or up to 1'000%. For the previously cured and swollen hydrogels, parallel plate with 10 mm diameter was used and the gap was set to have a normal force of around 0.1 N.

*hMSC Cell Culture*

*Expansion and Passaging*

Frozen cells were thawed rapidly at 37°C and suspended in Dulbecco's Modified Eagle's medium (DMEM) at 4°C. While centrifuging the cells at 300g for 10 min at 4°C, 90 mL expansion medium (DMEM with 10% fetal bovine serum (FBS), 1% antibiotic-antimycotic

(Anti-Anti), 1% non-essential amino acids and 0.001% bFGF aliquot) was loaded per triple flask. 10 mL of the cell suspension was then added to the flask. Incubation was done at 37°C with 5% $CO_2$. On day 3 (d3), 50 mL expansion medium was added and on d5, the medium was changed with 100 mL per flask. The cells were split when they reached 80% confluence. First, the cells were washed twice with PBS at 37°C. Then, 15 mL of 0.05% trypsin-EDTA was loaded. After incubation at 37°C for 4 min and tapping the flask to detach all cells, 15 mL control medium (DMEM with 10% FBS and 1% Anti-Anti) was added per flask and the cell suspension transferred to a falcon tube. After an additional washing step with 15 mL control medium, the cells were again centrifuged at 10 min for 10 min at 4°C. The cells were then resuspended in 10 mL control medium at 4°C and counted using a hemocytometer and trypan blue. Finally, the cells were resuspended at the desired concentration after another centrifugation step in expansion medium and loaded into a fresh cell culture flask. Medium was changed three times a week thereafter and the cells again split once confluence was reached.

*Live/Dead Assay*

After washing with warm PBS, a staining solution containing 1 mM Calcein AM and ethidium homodimer-1 (EthD-1) was loaded. The sample was then incubated for 15 min and washed twice with PBS. With the sample in PBS, the dish was sealed with parafilm and imaged within one hour. After that, PBS was replaced with osteogenic medium (control medium, 50 µg/ml ascorbic acid, 100 nM dexamethasone, 10 mM beta-glycerophosphate).

*Fixation and Actin-Nuclei Staining*

The samples were fixed with 4% PFA by washing with warm PBS, incubating with 4% PFA for 15 min at RT and washing twice with PBS. PBS was then added for hydration and the sample stored at 4 °C until the staining procedure was started. For the staining, the sample was first blocked with 1% BSA for 1 h and then permeabilized in 0.2% Triton X-100. After washing thrice with PBS, a staining solution was added, which contained 0.1% BSA, Phalloidin 647 and Hoechst (1 mg/mL) at dilutions of 1:200 and 1:1000 respectively. After 1-2 hours of incubation at RT protected from light, the sample was washed thrice with PBS and PBS was loaded for hydration.

*Cell Area Quantification*

For the cell area quantification, maximum intensity projections (MIPs) of three z-stacks were analyzed per group. The threshold was manually set and kept the same for all groups within one experiment. The area was measured by the actin signal. The cell nuclei were then manually counted. The average cell area was then calculated by the total area divided by the cell number for each z-stack.

*Alizarin Red S Staining*

Fixed volumetrically printed samples were stained for qualitative assessment of mineralization by Alizarin Red S (ARS). An ARS staining solution was prepared at 2 mg/mL in milliQ water, pH 4.3. Samples were stained for 15 minutes, washed thrice with PBS and stored in PBS for hydration.

*Imaging and Image Analysis*

The confocal microscopes Leica SP8 MP, Leica SP8 AOBS CARS and Zeiss LSM 880 Airyscan were used for imaging. For imaging the whole printed constructs, a stereomicroscope (Leica Stereo) was used. Microscopes were used with the software LAS X (Leica) or ZEN 2012 (Zeiss). Images and videos during the volumetric printing process were captured with the integrated camera of the volumetric printer. Image analysis was performed in Fiji.

### Supplementary Figures

### Figure S1

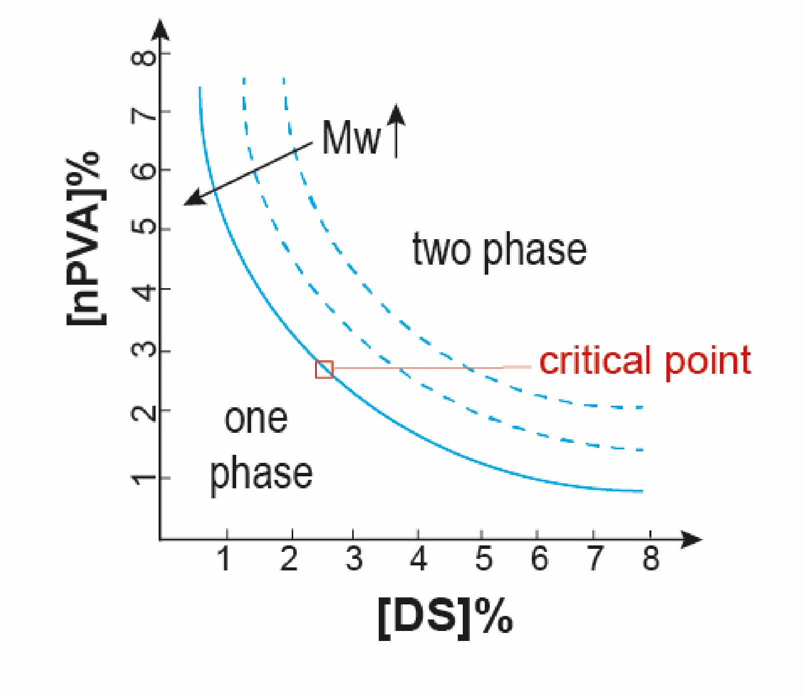


**Figure S1:** Schematic showing the phase diagram between nPVA and DS.

**Figure S2**

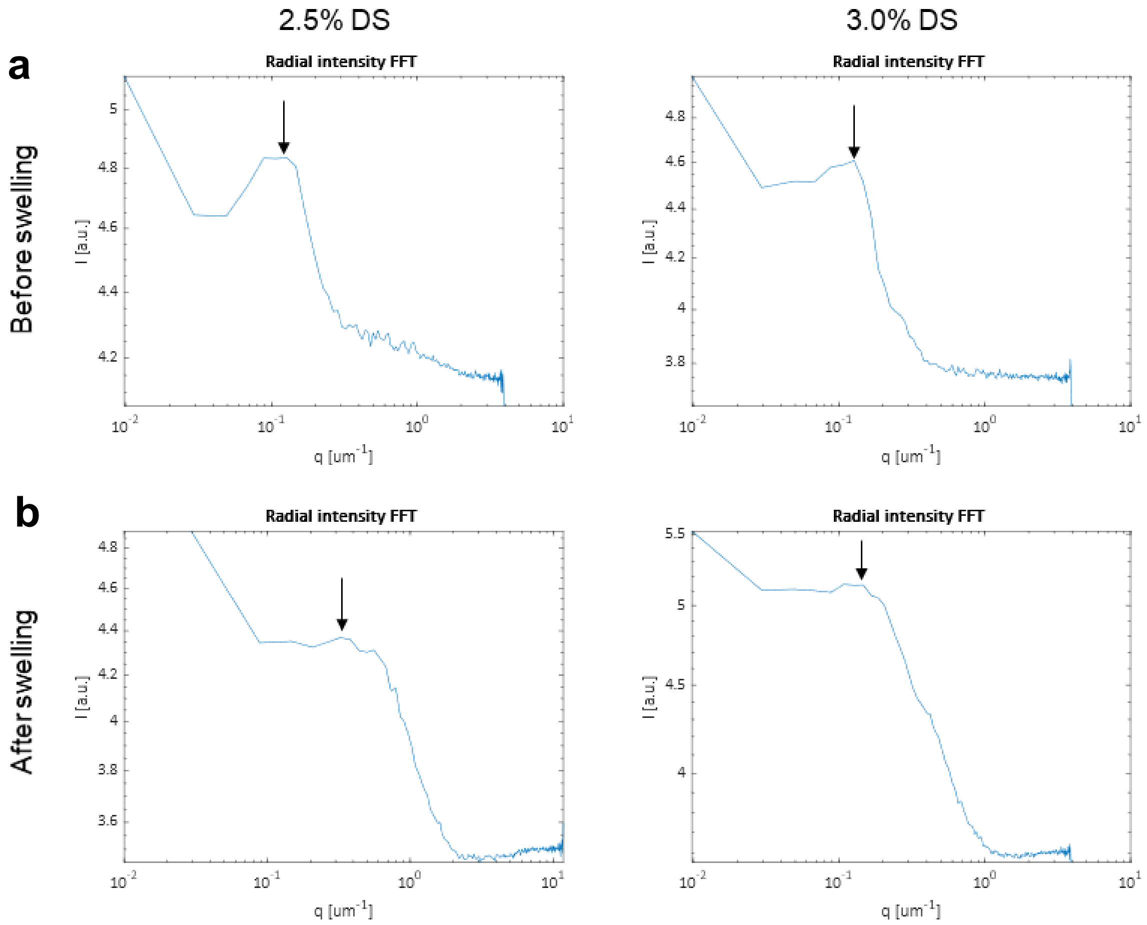


**Figure S2: RIFFT plots of macroporous hydrogels.** Hydrogels after UV crosslinking show a peak (arrow) corresponding to a length scale of 8 µm in Mix 1-2 before swelling (**a**) and 3 and 8 µm respectively in Mix 1-2 after incubation with FITC-dextran labelled pores (**b**).

### Figure S3

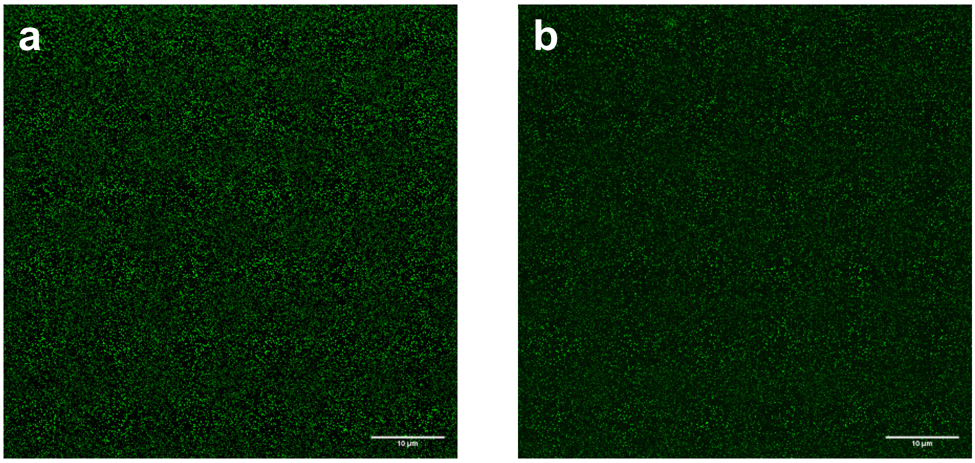


**Figure S3: Hydrogels with lower DS concentrations showing no pores.** 2% (w/w) nPVA (61 kDa, FITC-labelled nPVA : unlabelled nPVA = 1:50), with PEG-2-SH as crosslinker (thiol/ene ratio = 0.8); crosslinked with UV lamp (LAMAG); 2% (**a**) and 0% (**b**) (w/w) DS; unwashed; scale bars = 10 µm.

### Figure S4

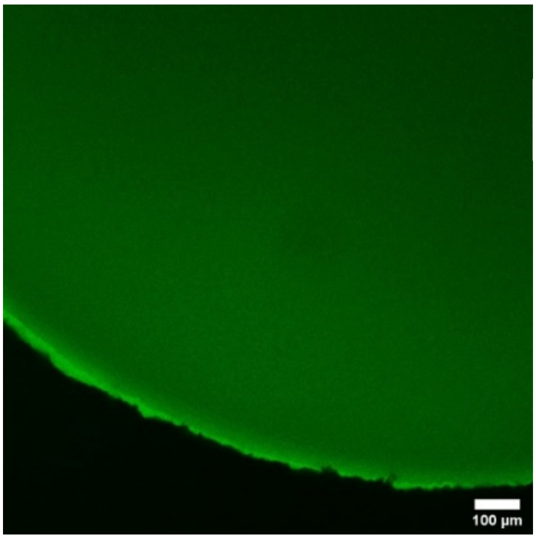


**Figure S4: Control gel without DS showing no pores**. CLSM images of 2% nPVA gels without DS after 3 days of equilibration in PBS and 2 days in FITC-dextran showing no pores.

**Figure S5**

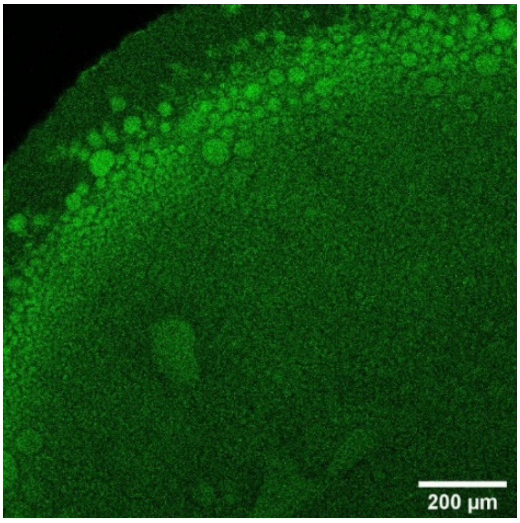


**Figure S5: Inhomogenously sized blobs indicating phase separation before photopolymerization with 3% nPVA 3.5% DS.** After UV curing (18 mW $cm^{-2}$, 365 nm), a 1/50$^{th}$ of FITC-labelled nPVA (green) was used for confocal imaging.

### Figure S6

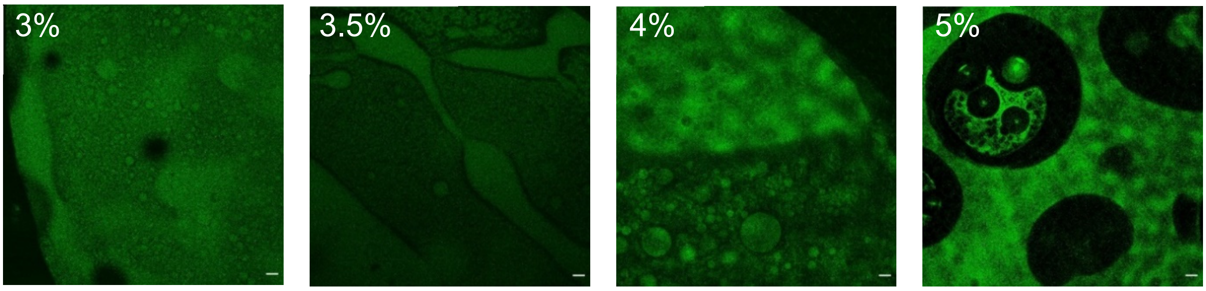


**Figure S6: Hydrogels with higher nPVA content showing inhomogeneous morphologies**. CLSM images of hydrogels with different concentrations of nPVA (47 kDa, 1:50 FITC-labelled nPVA : unlabelled nPVA), with PEG-2-SH as crosslinker (thiol/ene ratio = 1.04); crosslinked with UV lamp; 2.5% (w/w) DS; unwashed; 3%, 3.5%, 4%, 5% (w/w) nPVA. Scale bars = 50 µm.

### Figure S7

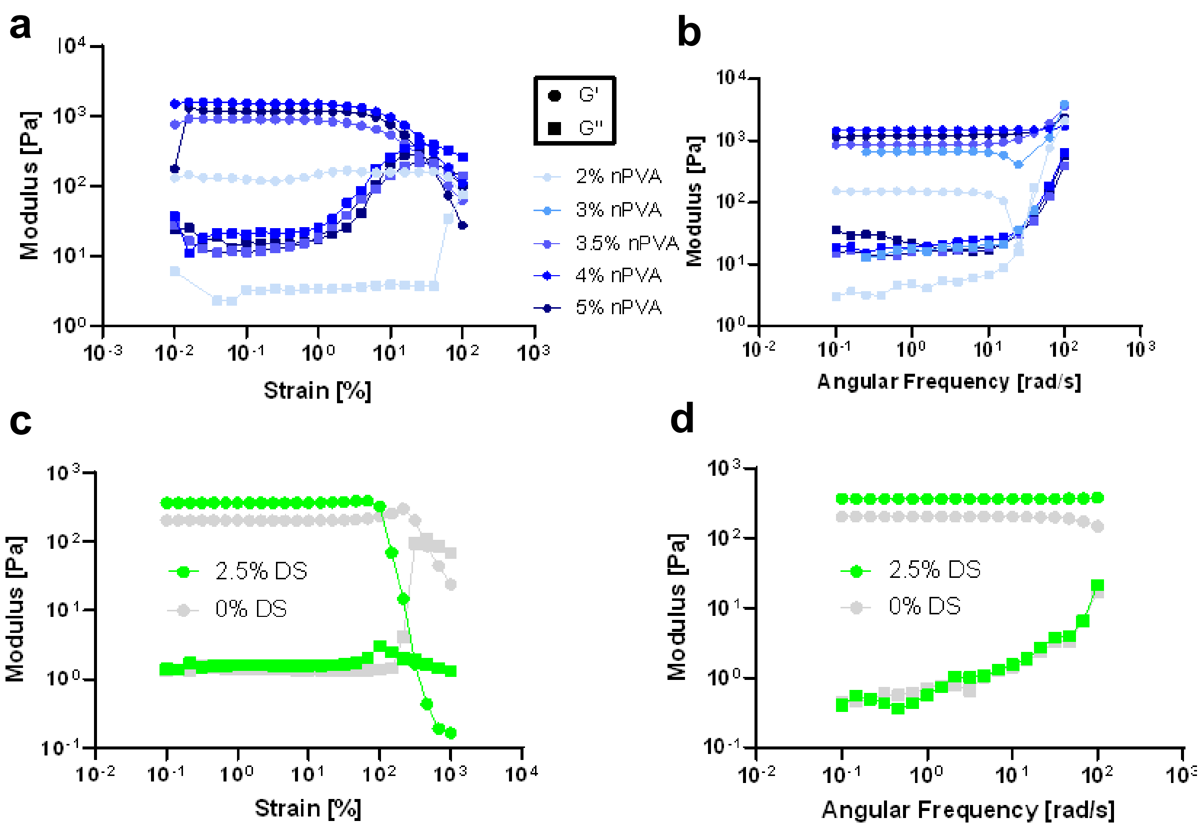


**Figure S7: Amplitude and frequency sweeps of various nPVA-DS hydrogels after photo-rheometry. a,b** The amplitude (**a**) and frequency (**b**) plots of hydrogel samples formed with varying nPVA% and 2% DS. **c, d** The amplitude (**c**) and frequency (**d**) plots of hydrogel samples formed with 0 vs. 2.5% DS and 2% nPVA. PEG-2-SH was used as a crosslinker (thiol/ene = 0.8). Amplitude sweeps: 10 rad/s angular frequency and 0.01-100% strain; frequency sweeps: 0.5% strain and 0.1-100 rad/s angular frequency.

**Figure S8**

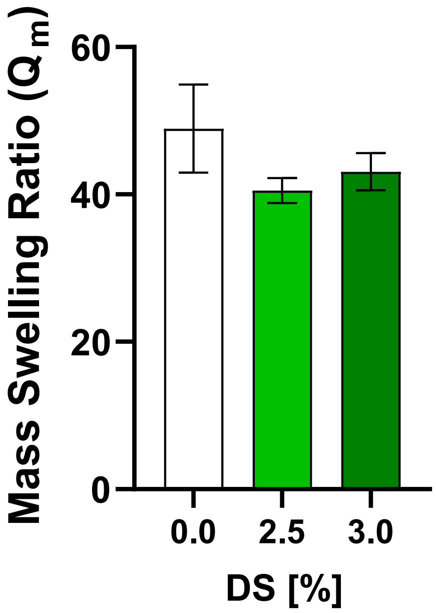


**Figure S8:** Equilibrium mass swelling ratio of hydrogels formed with 2% nPVA and varying DS content after two days of swelling in PBS.

**Figure S9**

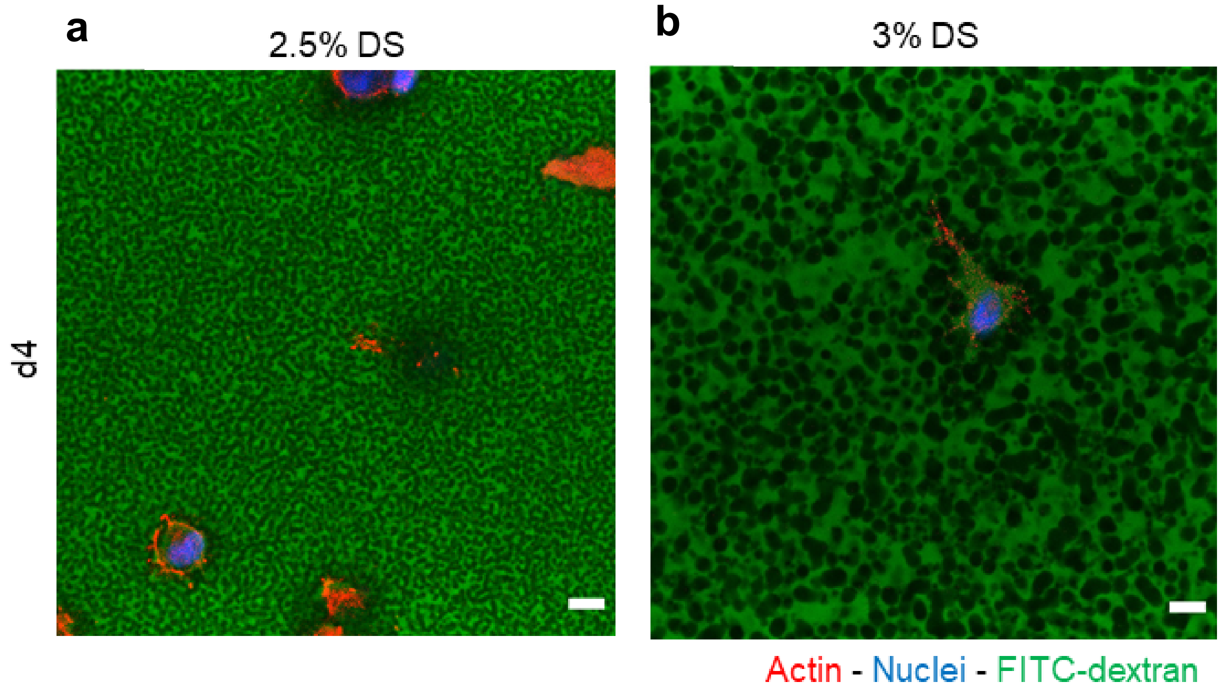


**Figure S9: Cell growth within macroporous nPVA hydrogels with different composition and porous microarchitecture.** Confocal images of gels labelled with FITC-dextran (green) showing smaller pores for 2.5% DS (**a**) and larger pores for 3% DS (**b**) with spreading hMSCs showing how cells penetrate through the pores. Scale bars: 20 µm.

### Figure S10

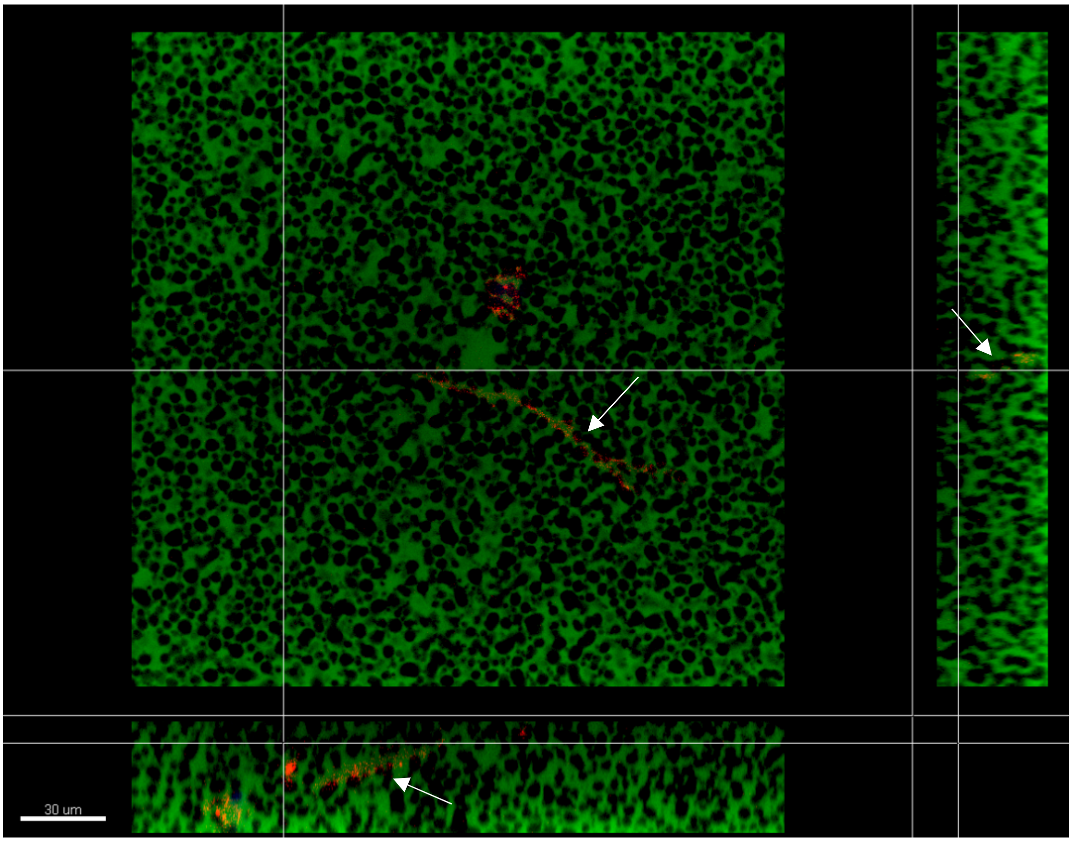


**Figure S10: Pores are penetrated by cell processes.** Orthogonally sectioned confocal z-stack taken showing how cells penetrate the pores (arrows) within a hydrogel, 2% nPVA with RGD motifs and PEG-2-SH crosslinker with 3% DS cured under UV (20 mW $cm^{-2}$, 365 nm). Hoechst (blue) and phalloidin 647 (red), FITC-dextran labelled pores (green).

**Figure S11**

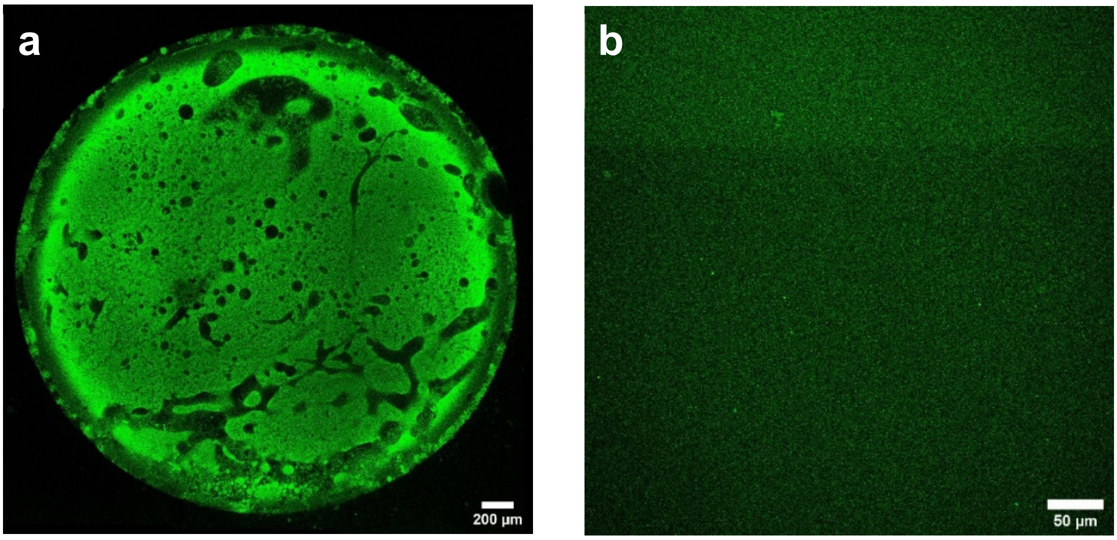


**Figure S11: 1.5% DS composition resulting in inhomogeneous pores and 0.5% DS composition resulting in no pores.** Confocal images of 2% nPVA 5% gelatin gels after washing with 1.5% DS showing large scale phase separation (**a**), 0.5% DS showing no phase separation at this resolution (**b**); a 1/50$^{th}$ of FITC-labelled nPVA (green) was used for confocal imaging.

**Figure S12**

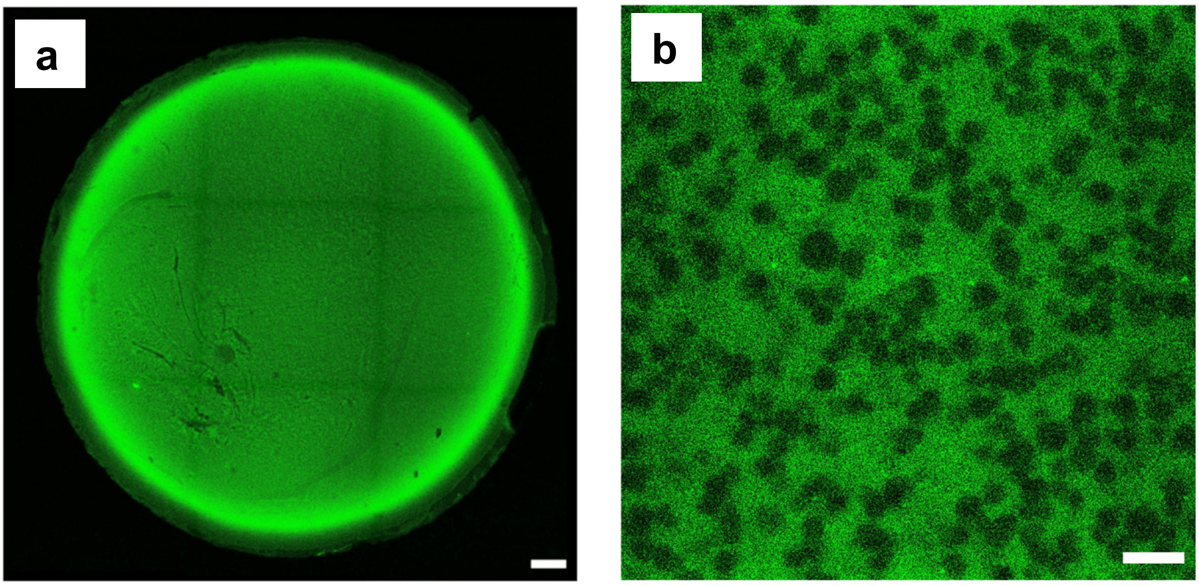


**Figure S12: Pore imaging using the optimized composition with gelatin.** Representative CLSM images of 2% nPVA (DoF = 20%) 5% gelatin 1% DS gel after UV curing (18 mW $cm^{-2}$, 365 nm) in cold state and incubating in PBS at 37°C for over one hour to release gelatin. A $1/50^{th}$ of FITC-labelled nPVA (green) was used for confocal imaging. **a** Whole well with almost complete homogeneity (merged image). **b** Gel at higher magnification within a homogenous region showing macropores. Scale bars: 200 µm (**a**) and 10 µm (**b**).

### Figure S13

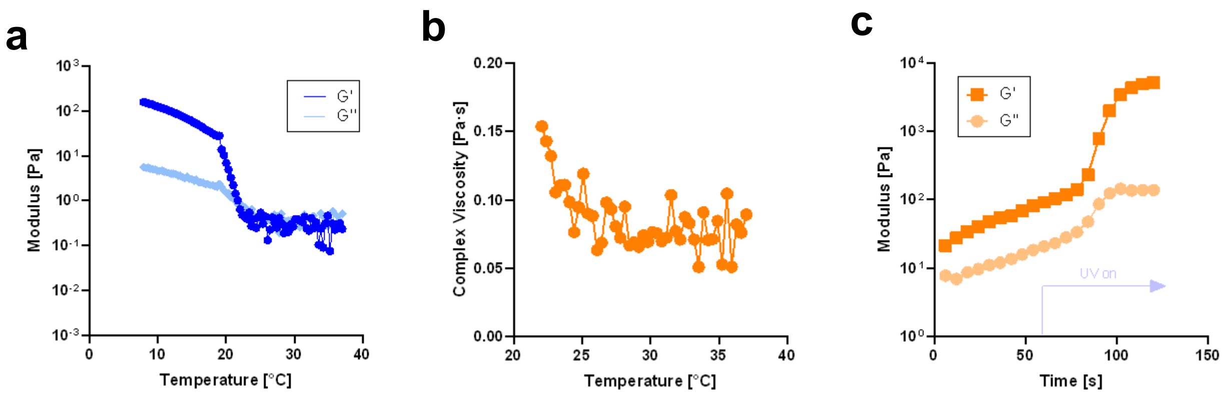


**Figure S13: Temperature-dependent gelation, complex viscosity and crosslinking of the resin. a** Temperature sweep of storage (G') and loss (G'') moduli from 37 °C to 8 °C showing the temperature-dependent gelation of gelatin at around 20-22 °C. **b** Complex viscosity obtained from the same measurements. **c** Time sweep of G' and G'' at 25 °C showing crosslinking after UV light (7.9 mW $cm^{-2}$, 365 nm) is turned on after 60 s of measuring. 2% nPVA (DoF = 20%) 1% DS 5% gelatin, drying prevented by mineral oil or wet tissue. 1 Hz, 0.5% (**c**) or 1% (**a, b**) strain, representative curves of 3 separate photo-rheometry experiments (N=3) shown.

### Figure S14

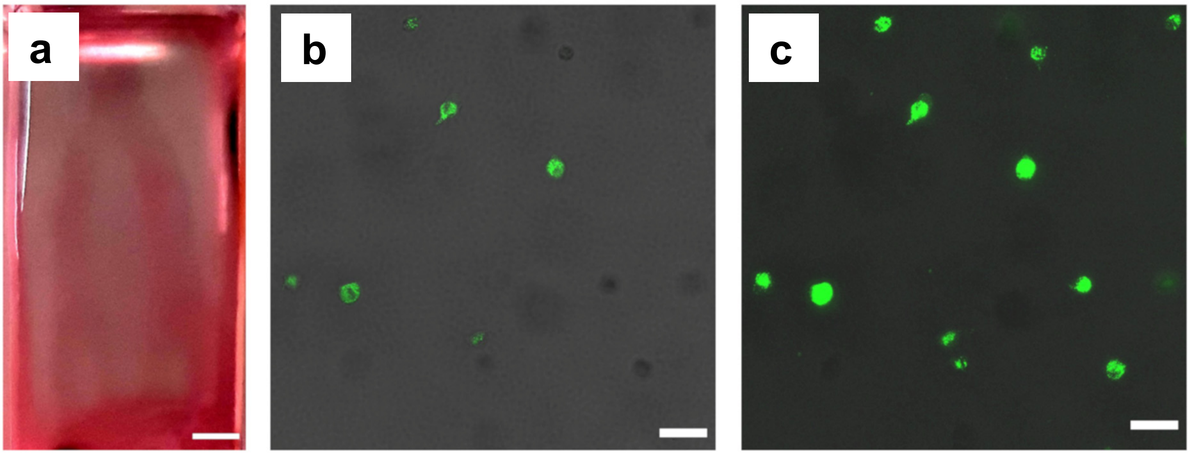


**Figure S14: Volumetric bioprinting of phase-separating resin**. **a** Image of the bioprinted branch structure taken after loading medium (red, contrast enhanced for better visibility). **b-c** Representative CLSM images of hMSCs stained with Calcein AM (green) showing alive cells, transmission channel (grey) showing pores in **b**; viability seems high considering the number of cells in focus the total amount of cells; note that some cells that are seen in the images are out of focus, which can be seen well when comparing **b** (single plane) and **c** MIP of 50 µm thick z-stack. Scale bars: 2 mm (**a**) and 50 µm (**b**,**c**).

**Figure S15**

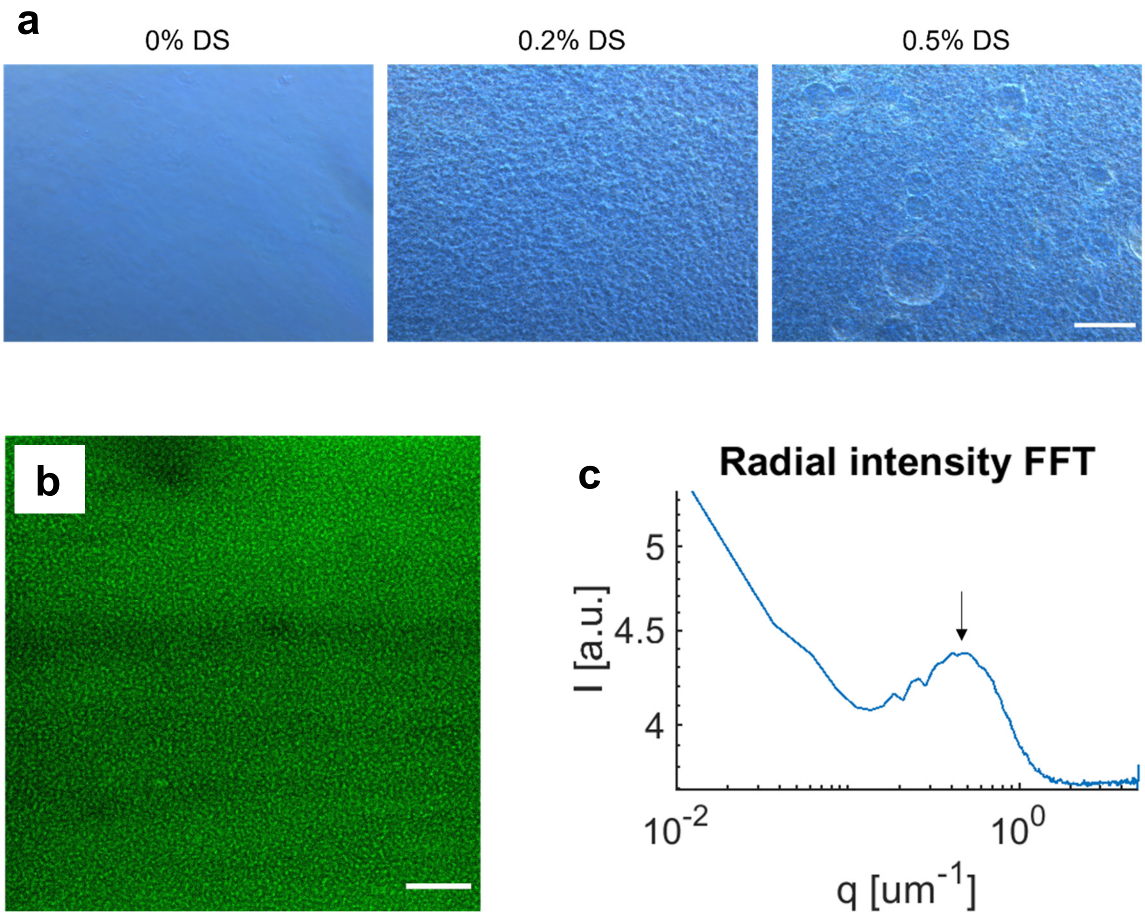


**Figure S15: Porosity in the adapted phase-separating bioresin after casting and volumetric bioprinting. a** Phase contrast microscopy images immediately after acellular gel casting of the adapted bioresin for VBP with varying amounts of DS. Scale bar = 100 µm. **b** CLSM images of a volumetrically bioprinted 2% nPVA, 0.2% DS, 4% gelatin hydrogel construct containing hMSCs, fixed at day 2 of osteogenic culture and stained for 1 day in FITC-dextran (green). **c** RIFFT of the image in **b** showing a peak corresponding to a length scale of 2.2 µm. Scale bar = 20 µm.

**Figure S16**

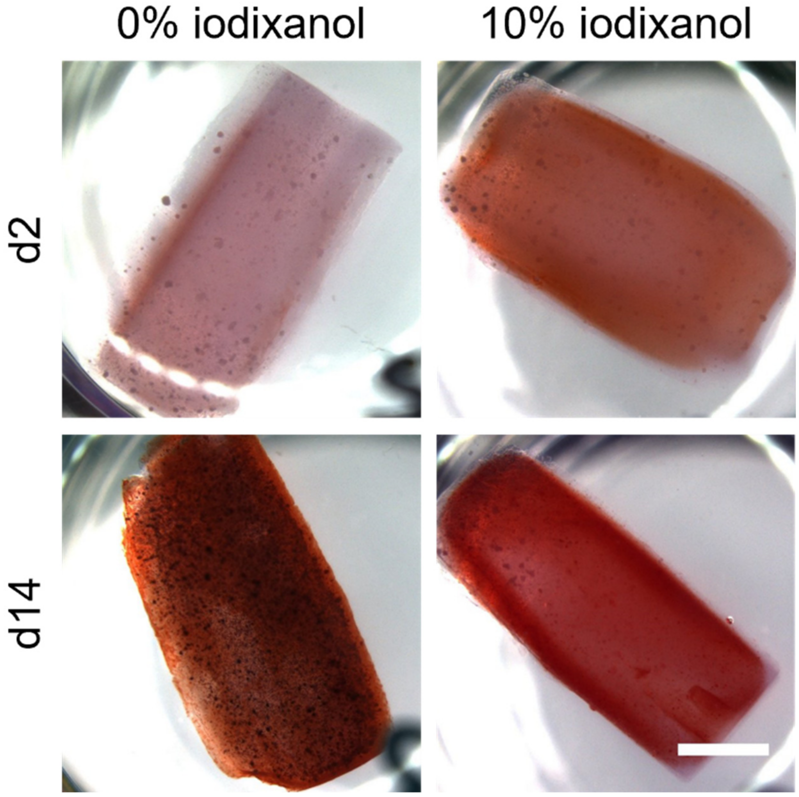


**Figure S16: Alizarin Red S staining of volumetrically bioprinted resins.** Using an adapted phase-separating resin, printed hMSC-laden constructs fixed at days 2 and 14 of 3D culture were stained with Alizarin Red S for qualitative analysis of mineralization. Scale bar: 2 mm.

**Figure S17**

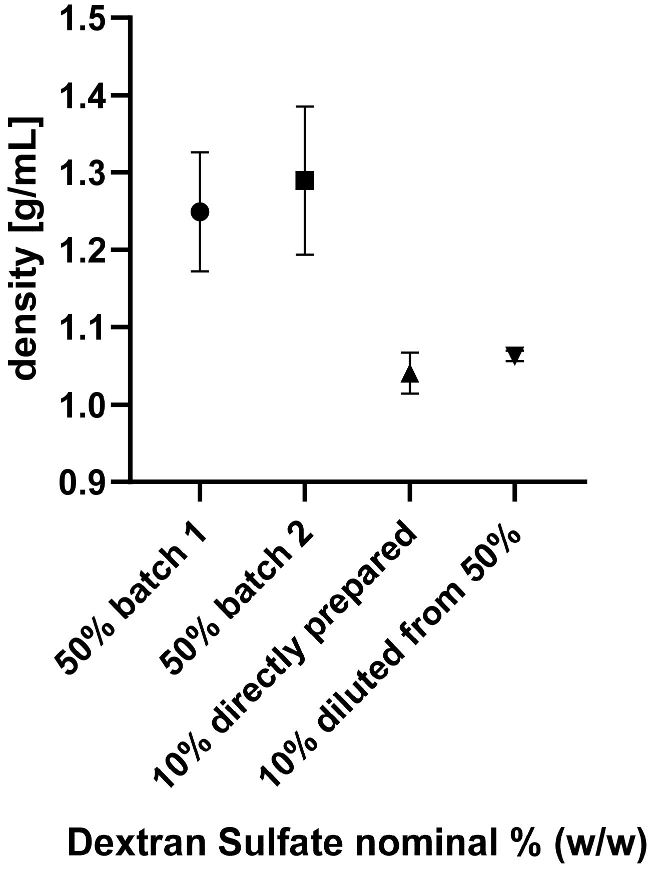


**Figure S17: Density of different DS stock solutions.** Obtained from weighing 10 µL, N=3, data presented as mean ± SD.

**Table S1: Results of the second laser dose test with 2% nPVA.** DoF = 20% with 5% gelatin and 1% DS, green = derived threshold 75 mJ cm$^{-2}$, less pronounced dots marked in grey.

| | Time [s] | | | | | | | |
|---|---|---|---|---|---|---|---|---|
| Intensity [mW cm$^{-2}$] | 44 | 39 | 34 | 29 | 24 | 19 | 14 | 9 |
| 6.6 | 290 | 257 | 224 | 191 | 158 | 125 | 92 | 59 |
| 5.8 | 255 | 226 | 197 | 168 | 139 | 110 | 81 | 52 |
| 5 | 220 | 195 | 170 | 145 | 120 | 95 | 70 | |
| 4.2 | 185 | 164 | 143 | 122 | 101 | 80 | | |
| 3.4 | 150 | 133 | 116 | 99 | 82 | 65 | | |
| 2.6 | 114 | 101 | 88 | 75 | 62 | | | |
| 1.8 | 79 | 70 | 61 | | | | | |
| 1 | 44 | | | | | | | |

**Table S2**: **Components of the 2% nPVA 2.5% DS composition.**

| **Component** | **Diluted in** | **Concentration [w/w]** | **Relative volume [v/v]** | **Final concentration [w/v]** |
|---|---|---|---|---|
| **nPVA (DoF =7%)** | 0.1 % LAP | 4% | 50% | 2% (0.05% LAP) |
| **DS** | PBS | “10%” | 25% | 2.5% |
| **PBS** | | 1x | 12.5% | |
| **PEG-2-SH (2 kDa)** | PBS | 15.34% | 12.5% | 1.9% (thiol/ene ratio: 0.8) |

**Table S3: Components of the 2% nPVA 3% DS composition used for cell embedding.**

| **Component** | **Diluted in** | **Concentration [w/w]** | **Relative volume [v/v]** | **Final concentration [w/v]** |
|---|---|---|---|---|
| **DS** | PBS | “10%” | 30% | 3% |
| **PBS** | | 1x | 8% | |
| **CGRGDSP** | PBS | 14.10% | 2% | 0.28% (10% of thiol groups) |
| **nPVA (DoF = 7%)**<br>**with cells** | 0.1 % LAP | 4%<br>1 Mio/mL | 50% | 2% (0.05% LAP)<br>0.5 Mio/mL |
| **PEG-2-SH (2 kDa)** | PBS | 16.09% | 10% | 1.9% (thiol/ene ratio: 0.8) |

**Table S4: Composition of the 2% nPVA 1% DS 5% gelatin resin for acellular volumetric printing.**

| Component | Diluted in | Concentration | Relative volume [v/v] | Final concentration [w/v] |
|---|---|---|---|---|
| **Gelatin** | PBS | 10% w/v | 50% | 5% |
| **nPVA (DoF = 20%)** | 0.25 % LAP | 10% w/w | 20% | 2% (0.05% LAP) |
| **PBS** | | 1x | 10% | |
| **DS** | PBS | 10% w/w | 10% | 1% |
| **PEG-2-SH (2 kDa)** | PBS | 16.09% w/w | 10% | 1.9% |

**Table S5: Composition of the 2% nPVA 0.2% DS 4% gelatin resin for volumetric bioprinting.**

| Component | Diluted in | Concentration | Relative volume [v/v] | Final concentration [w/v] |
|---|---|---|---|---|
| **nPVA (DoF = 20%)** | 0.25% LAP in PBS | 10% w/w | 20% | 2% (0.05% LAP) |
| **PBS** | | 1x | 17.3% | |
| **Pyrogallol** | PBS | 2.5 mg/mL | 1% | 25 mg/L |
| **CGRGDSP** | PBS, pH 6 | 20 mM | 5% | 1 mM |
| **DS** | PBS | 10% w/w | 2% | 0.2% |
| **Iodixanol** | OptiPrep | 60% w/v | 16.7% | 10% |
| **Gelatin**<br>**with cells** | PBS | 14.29% w/v<br>7.143 mio/mL | 28% | 4%<br>2 mio/mL |
| **PEG-2-SH** | PBS, pH 6 | 28.21% w/w | 10% | 3.78% (thiol/ene ratio: 0.8) |

Pyrogallol was added to prevent premature crosslinking. For the composition with 0% iodixanol, the iodixanol fraction was replaced with PBS.

**MATLAB Script for FFT**

```
%% FFT code

close all
clear
clc

%% Insert parameters and load image
% NB note that the image should be square!

im = imread('2021-03-05_E17.4_branch_95_pores_FITCdex_onlymerged.lif - 
25x_closetoborder.jpg');
conv = 1/1.65; %specify scalebar [Âµm/pxl] = 1/[px/um], ALWAYS CHECK IF 
TRUE 1/16.6320 (63x z3), 1/11.0880 (63xz2) 1/5.5440 (63x), 1/1.65 (25x),
1/1.7600 (20x),1/0.8800 (10x)

%% Automatic

grayImage = rgb2gray(im); %greyscale
figure
imagesc(grayImage);
grayImage=double(grayImage);  %NEW because error using  .* otherwise
[rows columns numberOfColorChannels] = size(grayImage);
if numberOfColorChannels > 1
    grayImage = rgb2gray(grayImage);
end

% Convert from pixel size to "future" q space in 2D
matsize = size(grayImage);
L = matsize(1); %Pixel size of your image. If this is a square, matsize(1) 
and matsize(2) are the same
LL=matsize(2);
FOV = L*conv; %[Âµm] field of view of the image
Qmax = (L/2)/FOV; % Maximum wave vector
Qx = linspace(-Qmax, Qmax, L);
Qy = linspace(-Qmax, Qmax, L);

wc=hamming(L); % Make a window
[maskr,maskc]=meshgrid(wc,wc);
w=maskr.*maskc; % This is the window
fftOriginal = fft2(double(grayImage.*w)); % Take the 2d FFT of the image 
multiplied by the window

shiftedFFT = fftshift(fftOriginal); % Shift so that the centre is at the 
centre of the image
[QQx,QQy] = meshgrid(Qx,Qy);
Qr=sqrt(QQx.^2+QQy.^2);

I.fft = log10(abs(shiftedFFT));

figure
imagesc(log10(abs(shiftedFFT)),'Xdata', Qx, 'Ydata',Qy);
axis('image')
xlabel('Qx [1/um]'); ylabel('Qy [1/um]');
set(gca,'FontSize',12,'FontName','Calibri Light');
set(gcf,'color','w')
title('FFT')

% Do the radial binning
k=0;
for i=1:size(Qr,2)
    for j=1:size(Qr,1)
```

```
        k=k+1;
        PR(1,k)=Qr(i,j);
        PR(2,k)=log10(abs(shiftedFFT(i,j)));
    end
end
[~,idx] = sort(PR(:,1)');
sortedPR = PR(idx,:)';
[RBin,PropertyBin]=radial_binning(PR(1,:)',PR(2,:)',200,max(PR(1,:)));

I.RBin = RBin;
I.PropertyBin = PropertyBin;

figure
plot(RBin,PropertyBin, 'LineWidth',1.5)
xlabel('q [um^-^1]');
ylabel('I [a.u.]');
title('Radial intensity FFT')
set(gca,'FontSize',24,'FontName','Arial');
set(gcf,'color','w')
set(gca,'Yscale','log','XScale','log')
set(gca,'box','off')
set(gca,'LineWidth',1.5)
```

---

```
function [RBin,PropertyBin]=radial_binning(R,I,BinNo,MaxR)

%% --------------------------------------------------------------------------
% Parameters
% --------------------------------------------------------------------------
% Parameters of Geometry:

% Parameters for Binning:
number_of_rings = BinNo;        % number of Bins for Binning

%% --------------------------------------------------------------------------
% Not-to-be-messed-with Parameters
% --------------------------------------------------------------------------
R_min=0;             % Minimal Distance from Center at which
% positions are considered
R_max = MaxR; % Maximal Distance from Center at which
% positions are considered

rmin = R_min;
rmax = R_max;

Bin_Edges = linspace(rmin,rmax,number_of_rings+1);  % Edges of Bins
Bin_Centers=(Bin_Edges(1:end-1)+Bin_Edges(2:end))./2; % Center of Bins.

RadialPos=R;
Property=I;

%% --------------------------------------------------------------------------
-
% Binning
%---------------------------------------------------------------------------
% The Particles are Binned according to the predefined Bins radially.

% Pre-initialize Matrices.
PropertyMean=[];

[n,whichbin] = histc(RadialPos,Bin_Edges);
for i=1:number_of_rings
    flagBinMembers=(whichbin==i);
```

```
    binMembers=Property(flagBinMembers);
    PropertyMean(i)=nanmean(binMembers);
end

%% ------------------------------------------------------------------------
% Rename for clarity and subtract Brownian Motion
%-------------------------------------------------------------------------
PropertyBin=PropertyMean;
RBin=Bin_Centers;

end
```

**Supplementary Movies**

**Movie 1 -** Animation of phase-separated hydrogels with FITC-dextran-labelled porous microstructure. Scale bar, 20 μm.

**Movie 2 -** Animation of actin-nuclei-stained hMSC cells in phase-separated hydrogels at day 13 following up in vitro osteogenic culture. Scale bar, 200 μm.

**Movie 3 -** Volumetric printing process of the resin supplemented with gelatin within 12 seconds.

**Movie 4 -** Volumetric bioprinting process within 18 seconds. The optically tuned bioresin contained hMSCs at a concentration of $2 \times 10^6$ cells/mL and 10% iodixanol.
